# Host background shapes the portability of a non-canonical translation initiation system across *Escherichia coli* strains

**DOI:** 10.64898/2026.04.16.719103

**Authors:** Dominic Scopelliti, Andras Hutvagner, Paul R. Jaschke

**Author notes:** To whom correspondence should be addressed: Paul R. Jaschke. These authors contributed equally (Dominic Scopelliti, Andras Hutvagner).

## Abstract

Translation initiation has become an attractive target for engineering orthogonal translation systems, yet the extent to which these systems retain functionality across distinct host backgrounds remains poorly defined. In bacteria, start codon recognition depends on pairing between the initiator tRNA anticodon and a suitable start codon within the appropriate distance from the Shine-Dalgarno sequence. These sequence-specific interactions enable translation initiation to be reprogrammed through anticodon engineering. What is currently missing is an understanding of how anticodon mutants of initiator tRNAs function across different bacterial strains. Here, we systematically evaluated the portability of a library of twelve i-tRNA anticodon mutants paired with their complementary non-canonical start codons. Most i-tRNA–start codon pairs supported detectable translation initiation across multiple strains, demonstrating broad functional portability. However, initiation efficiency, absolute system output, and fitness effects varied substantially between strains. Comparative genomic analyses revealed host-specific gene differences broadly, and endogenous tRNA gene sequence and copy number specifically, was associated with this variability. While most i-tRNA variants were well tolerated, a subset produced strain-dependent growth defects that primarily affected growth rate rather than final culture density. Together, these findings show that translation initiation efficacy of engineered i-tRNAs is partially strain-dependent and that host background must be considered a key design variable when deploying these translation systems. Looking forward, this study provides a framework for host-aware selection of microbial chassis for orthogonal translation applications in synthetic biology.

**Graphical Abstract:** 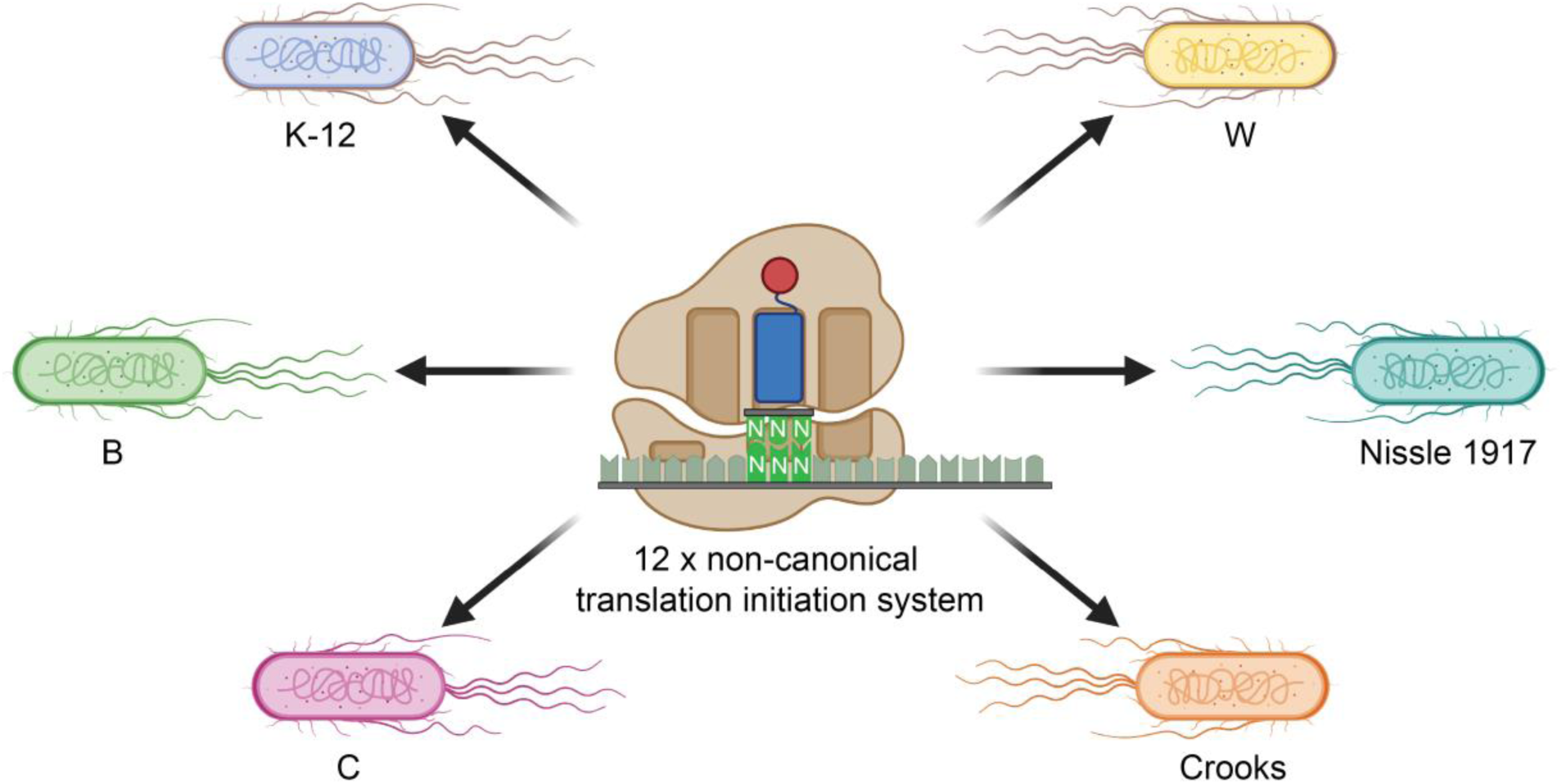

## Introduction

The deployment of orthogonal translation systems in synthetic biology has increasingly enabled precise control over protein expression, expansion of the genetic code, and the introduction of novel regulatory layers into engineered biological systems (*1*). By establishing independent channels of gene expression that operate orthogonally to native cellular machinery, these systems can reduce competition for translational resources, improve the performance of synthetic gene circuits, establish biocontainment models, and enable the incorporation of non-canonical amino acids into proteins (*2–5*). As a result, orthogonal translation has become a powerful strategy for programming cell behavior and expanding the functional capabilities of engineered organisms (*1*). One promising strategy for designing orthogonal translation systems involves engineering translation initiation, the primary regulatory checkpoint of protein synthesis, where start codon recognition occurs on messenger RNA (mRNA) transcripts (*6, 7*). As recognition is primarily mediated through interactions between the ribosome with the Shine-Dalgarno (SD) sequence and the anticodon on initiator tRNA (i-tRNA) with the mRNA start codon, modifying the i-tRNA interaction provides an attractive approach for redirecting translation initiation to alternative non-canonical codons while minimizing interference with native translation processes.

In bacteria, translation initiation requires the coordinated assembly of the i-tRNA, mRNA, both the 30S and 50S ribosomal subunits, and initiation factors (IFs) 1–3 to form the mature 70S initiation complex (70S IC)(*6, 8–10*). During SD-mediated initiation, the 30S ribosomal subunit recruits mRNA through base-paring between the SD sequence in the 5’ untranslated region and the anti-SD sequence on the 16S rRNA, while the i-tRNA CAU anticodon recognizes the AUG start codon (*11*). This i-tRNA—start codon interaction is a critical regulatory checkpoint, inducing conformational changes that stabilize the 30S pre-initiation complex and drive its irreversible transition to form the elongation-competent 70S IC (*7, 11–13*). Several comprehensive reviews of bacterial translation initiation mechanisms are available in the literature (*6, 7, 13, 14*).

This dependence on i-tRNA—start codon base-pairing has enabled the development of engineered i-tRNAs capable of recognizing non-canonical start codons. By mutating the anticodon while preserving the conserved structural motifs that distinguish i-tRNAs from elongator tRNAs, synthetic i-tRNAs that selectively initiate translation from complementary non-canonical start codons have been developed (*15–19*). These systems have also been used in conjunction with engineered aminoacyl-tRNA synthetases to incorporate non-canonical amino acids at the N-terminus of proteins, expanding the chemical diversity of target proteins (*20–22*). For example, wild-type and mutant tyrosyl-tRNA synthetase from *Methanocaldococcus jannaschii* and Pyrrolysyl-tRNA synthetase from *Methanosarcina barkeri or mazei* has enabled the incorporation of more than 150 non-canonical amino acid analogues at the N-terminus of recombinant proteins (*21*).

However, one aspect that remains poorly characterized is the portability of these systems across different hosts. Engineered genetic systems that function robustly in one strain can exhibit unpredictable behavior in another, owing to differences in cellular physiology, gene regulation, tRNA abundance, and translational capacity (*23*). Such variability in orthogonal translation system performance could significantly constrain the reliability of synthetic gene circuits when deployed across different hosts – a growing concern as synthetic biology increasingly shifts toward the development of specialized microbial chassis for industrial biotechnology and therapeutic applications. Addressing this limitation requires a systematic evaluation of both the portability and biological tolerance of engineered i-tRNAs across multiple strains, ultimately enabling the development of design principles for robust orthogonal translation system performance.

*E. coli* provides an ideal model system to investigate this phenomenon, as many generally regarded as safe laboratory (GRAS) strains with distinct genetic backgrounds are already widely used in the production of industrially valuable compounds and pharmaceuticals (*24, 25*). Strains such as K-12, Crooks, W, B, and C are widely used in industrial biotechnology for the production of compounds such as beta-carotene, vitamin B12, succinate, ethanol, and itaconic acid (*26–32*). The probiotic strain Nissle 1917 has been clinically used for the treatment of intestinal disorders such as ulcerative colitis and inflammatory bowel disease (*33–37*). More recently, advances in synthetic biology have enabled the engineering of Nissle 1917 as a platform for biotherapeutics (*38–43*), as well as focused efforts on improving its potential as a chassis organism (*44, 45*). However, because these strains exhibit substantial differences in genome composition, regulatory networks, and translational capacity, the efficiency and biological tolerability of engineered translation systems may vary significantly across hosts and ultimately impact their utility as microbial production platforms (*46*).

To address this gap, we investigated whether the performance of an engineered translation initiation system is conserved across diverse *E. coli* host backgrounds or instead shaped by chassis-specific translational context. Using a library of sfGFP reporters bearing complementary non-canonical start codons, we quantified the activity of 12 mutant i-tRNAs across six commonly used *E. coli* strains and assessed both system output and host fitness consequences. We show that non-canonical initiation is broadly functional but not uniformly portable, where inducibility, absolute expression output, and physiological tolerance all vary substantially across strains and i-tRNA—start codon pairs. Comparative genomic analyses further identify candidate host features that may contribute to these differences. Together, this work establishes host background as an important design variable in the deployment of non-canonical translation initiation systems and provides a framework for selecting strain—system combinations suited to different synthetic biology applications.

## Results and Discussion

### Strain selection and design of a non-canonical initiation system for cross-strain analysis

To determine whether non-canonical translation initiation behaves as a portable or a chassis-dependent property, we compared the performance of 12 engineered i-tRNAs (Figure 1A) across a panel of genetically diverse and commonly used *E. coli* strains. To guide strain selection, we expanded on previous work characterizing an amber i-tRNA (*47*). Specifically, we constructed a phylogenetic tree of 18 common *E. coli* strains (Figure 1B). From this analysis, six strains were identified as phylogenetically distinct and widely used across research, industrial biotechnology, and clinical applications. These included the laboratory strains K-12 (MG1655), B (BL21(DE3)), C (C122), Crooks (ATCC 8739), and W (ATCC 9637), which are generally regarded as safe and classified as risk group 1 organisms, as well as the probiotic strain Nissle 1917. Selection of phylogenetically distinct strains allowed us to evaluate how differences in cellular physiology and translational environments influence the performance of non-canonical translation initiation systems.

**Figure 1.**
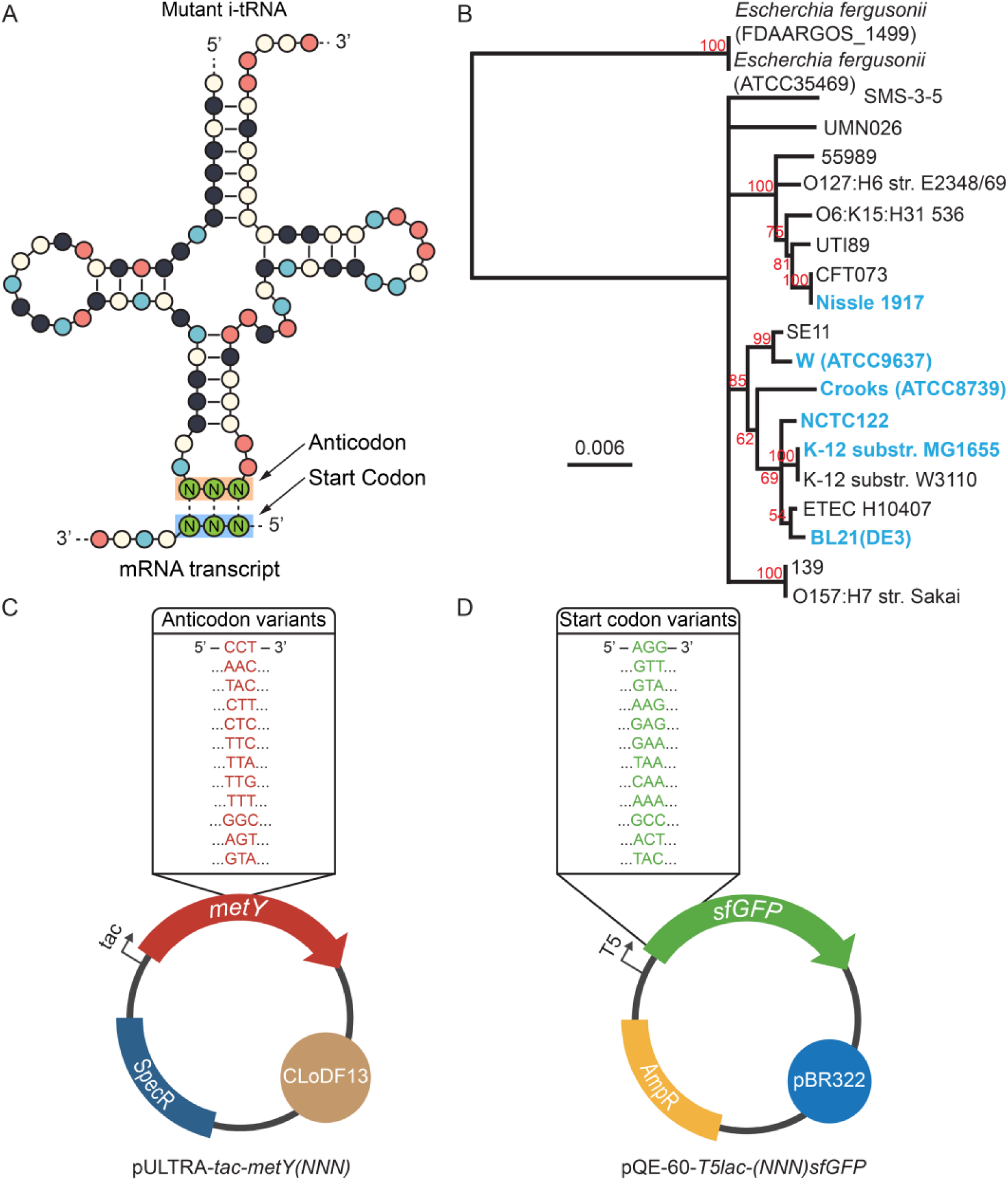
Design and experimental framework for evaluating non-canonical translation initiation systems. **(A)** i-tRNA secondary cloverleaf structure with altered anticodon region bound to a non-canonical start codon. Nucleotide identity indicated by color: (red) adenine, (yellow) cytosine, (light blue) uracil and (dark blue) guanine. Green colored nucleotides highlight the i-tRNA—start codon interaction. **(B)** Phylogenetic tree of 18 different strains of *Escherichia coli* with *Escherichia fergusonii* as an outgroup. Analysis was based on a MUSCLE multiple sequence alignment of concatenated multisequence locus typing (MLST) gene sequences *adk*, *recA*, *purA*, *fumC*, *gyrB*, *mdh*, *icd*. *E. coli* strains selected for this study are shown in blue. Consensus support (%) values are shown in red. Scale bar indicates the number of nucleotide substitutions per site. **(C)** Set of pULTRA-*tac-metY* plasmids containing medium copy CLoDF13 origin of replication, bearing a spectinomycin resistance (SpecR) gene and variants of the *metY* gene with altered anticodon sequences controlled by a *tac* promoter. **(D)** Set of pQE-60-*T5lac-sfGFP* medium copy plasmids with a ColE1 origin of replication, bearing an ampicillin resistance (AmpR) gene and variants of the *sfGFP* gene with altered start codons controlled by a T5-lac promoter.

To assess non-canonical translation initiation across these hosts, we generated a library consisting of 12 anticodon mutant i-tRNA plasmids paired with complementary sfGFP reporter constructs containing the corresponding start codons (Figure 1C). Both the i-tRNA and reporter plasmids were carefully designed to ensure compatibility of these systems across different strains. The i-tRNA anticodon mutants were cloned into a pULTRA plasmid and controlled by an inducible tac promoter (Figure 1D), while the sfGFP reporters were built on a pQE-60 backbone under the control of the T5-lac promoter, which is readily recognized by endogenous *E. coli* σ^70^ RNA polymerase, ensuring compatibility of the reporter system across the different strains. Each complementary pair of plasmids expressing the mutant i-tRNA and corresponding sfGFP reporter were then co-transformed into each of the six strains, generating a total of 72 strains for analysis. Bulk fluorescence from sfGFP expression was used as a measure of translation initiation efficiency across the different mutant i-tRNA—start codon pairs.

### Non-canonical translation initiation systems are functional across diverse E. coli strains

Having chosen our strains and plasmid designs, we first sought to assess whether engineered i-tRNAs support detectable non-canonical initiation across diverse *E. coli* hosts and whether the inducible dynamic range is consistent across backgrounds. To do this, we quantified reporter fluorescence for 12 mutant i-tRNA—start codon pairs across six *E. coli* strains under induced and repressed conditions. Analysis of fluorescence measurements revealed that non-canonical translation initiation was detectable in most strains, demonstrating that engineered i-tRNAs can function across diverse backgrounds (Figure 2). However, the magnitude of fluorescence varied substantially between both host strains and i-tRNA—start codon combinations, indicating that system output is strongly influenced by host-specific cellular contexts. To account for differences in baseline expression between strains and isolate system activity, fluorescence data were further analyzed as log_2_ fold changes (induced versus repressed conditions). A two-way ANOVA revealed significant effects on strain (p < 0.001), i-tRNA (p < 0.001), as well as strain and i-tRNA interaction (p < 0.001) (Table S1). This indicated that the efficiency of non-canonical initiation is highly dependent on both factors and their interaction, highlighting that individual i-tRNA—start codon pair behavior cannot be interpreted independently of the host translation environment.

**Figure 2.**
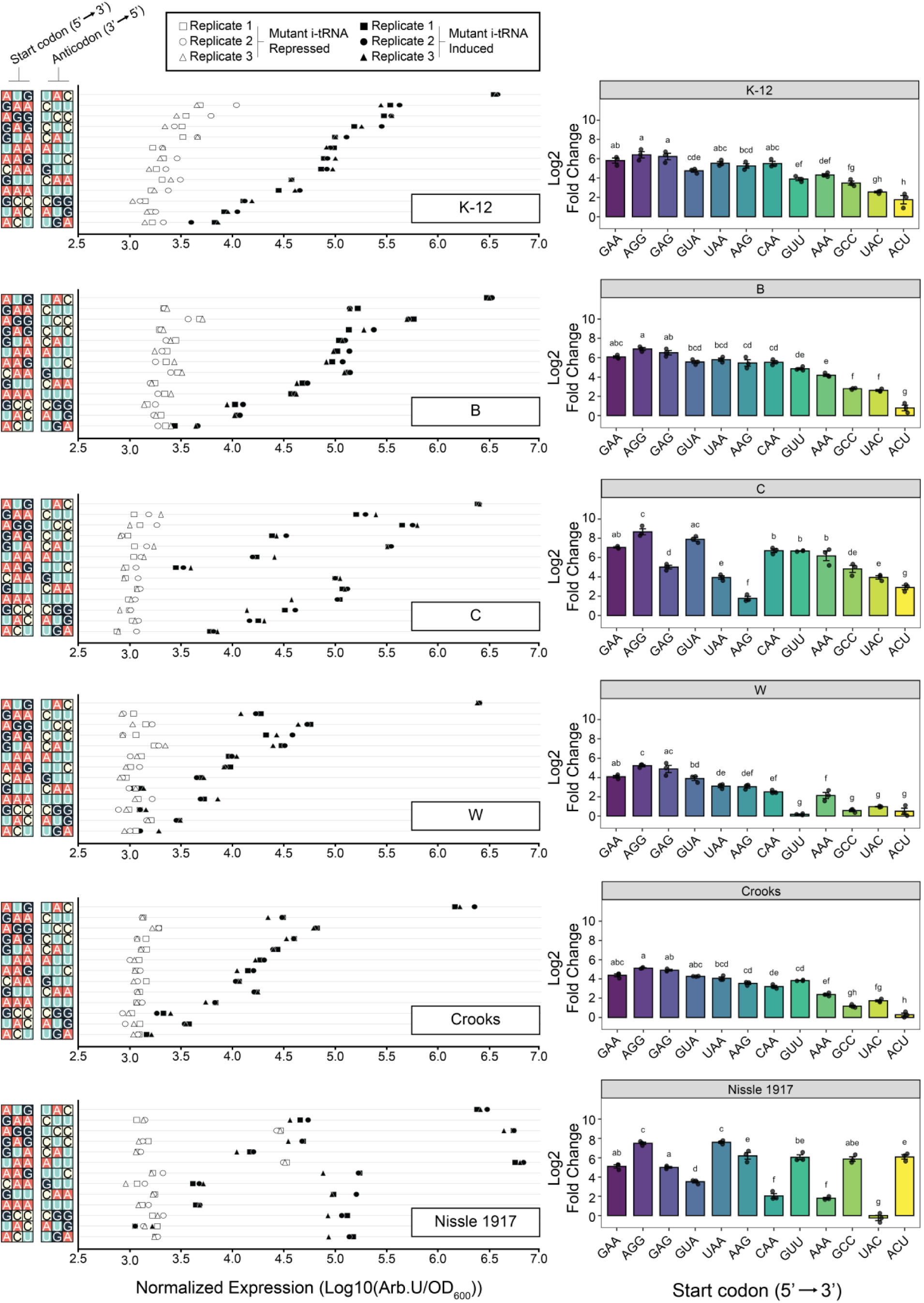
Non-canonical translation initiation across *E. coli* strains reveals strain and codon dependent differences in sfGFP reporter expression. (Left Panels) Normalized fluorescence levels (Arb. U./OD_600_) for each of the 12 engineered i-tRNA—start codon pairs across six *E. coli* strains (K-12, B, C, W, Crooks, and Nissle 1917) under repressed (open symbols) and induced (filled symbols) conditions. Cells were grown to mid-log phase (0.6 OD_600_), and induced cells (1 mM IPTG) were compared to control repressed cells (2% v/v glucose). Individual data points represent biological replicates (n=3). Fluorescence values reflect absolute reporter output and highlight differences in maximal expression capacity across strains. (Right panels) Log_2_ fold change values were calculated from paired OD_600_ normalized fluorescence measurements to account for differences in cell density and baseline expression. Bars represent standard error. Statistical analysis was performed using a two-way ANOVA followed by Tukey’s HSD post hoc testing. Compact lettering displayed above bars indicates statistically significant differences between codons within a given strain (p < 0.05), where letters that are common across codons indicate statistically indistinguishable groups.

Across all strains, several consistent i-tRNA—start codon trends were observed. The AGG start codon paired with the i-tRNA CCU repeatedly produced the largest fold-changes, whereas the ACU start codon paired with the i-tRNA-AGU showed little or no detectable fold-change in most strains. These trends suggest that certain i-tRNA—start codon combinations may be intrinsically more compatible with the bacterial initiation machinery, while others are poorly tolerated. Despite these shared patterns, substantial strain-dependent differences emerged, suggesting functionally relevant variation in translational machinery composition and capacity across hosts.

In the common laboratory strain K-12, several i-tRNAs exhibited strong activity, with an 80-fold increase for start codon AGG (i-tRNA-CCU), 76-fold increase for GAG (i-tRNA-CUC), and 52-fold increase for GAA (i-tRNA-UUC). However, analysis of raw fluorescence revealed elevated reporter expression under repressed conditions across all i-tRNAs (Figure S1 and Table S2). Because this increase was not restricted to specific codons, it may reflect global strain-specific effects rather than properties of individual i-tRNA-start codon pairs. One possible explanation is transcriptional leakiness of the mutant i-tRNA and sfGFP reporter, potentially arising from differences in the endogenous regulatory landscape of K-12. Alternatively, repression of this system relies on glucose-mediated catabolite repression, and strain-specific differences in glucose uptake or metabolism (*48*) could alter intracellular cAMP levels and reduce the effectiveness of CRP-cAMP-mediated repression. Such effects could contribute to incomplete transcriptional silencing and increased basal expression of the mutant i-tRNA and sfGFP reporter under nominally repressed conditions.

The widely used expression strain B, exemplified by BL21(DE3), also supported robust non-canonical initiation in our experiments. As observed in the K-12 strain, AGG (i-tRNA-CCU) produced the largest fold-change increase (118-fold), although background expression was slightly elevated compared to other strains (Figure S1). Notably, several additional pairs including, UAA (i-tRNA-UUA), AAG (i-tRNA-CUU), GAA (i-tRNA-UUC), GAG (i-tRNA-CUC), GUA (i-tRNA-UAC), and CAA (i-tRNA-UUG), produced similar fluorescence outputs. This reduced variability across multiple non-canonical i-tRNA–start codon pairs suggests a more uniform translation profile in this strain, which may be advantageous for applications requiring balanced expression from multiple engineered start codons. However, the slightly elevated baseline expression indicates that more stringent promoters, lower-copy plasmids, or genomic integration may be necessary for i-tRNA activity that functions similarly to canonical translation initiation.

In contrast to B strain, the Crooks strain exhibited comparatively low background fluorescence and more uniform expression under repressed conditions (Figure S1), indicating minimal leaky initiation. Upon induction, AGG (i-tRNA-CCU) produced the largest fold-change increase (35-fold), while several additional pairs including, GAA (i-tRNA-UUC), GUA (i-tRNA-UAC), UAA (i-tRNA-UUA), GUU (i-tRNA-AAC), AAG (i-tRNA-CUU), CAA (i-tRNA-UUG) and AAA (i-tRNA-UUU), generated moderate increases in translation (5—29-fold). This profile suggests that Crooks supports non-canonical initiation with relatively tight regulation but reduced overall efficiency, making it a potentially useful system for applications requiring low background expression and controlled activation.

Among the strains tested, the C strain exhibited the most permissive translational environment. All 12 tested i-tRNA—start codon pairs were functional in this strain, but produced a wide-range of translation levels. The most substantial increases were observed from AGG (i-tRNA-CCU) (402-fold increase), GUA (i-tRNA-UAC) (234-fold increase), GAA (i-tRNA-UUC) (130-fold increase), indicating that the C strain supports both high-efficiency initiation as well as broad compatibility across multiple i-tRNA–start codon pairs.

The W strain was among the least accommodative hosts. Although AGG (i-tRNA-CCU) again generated the largest fold-change increase (37-fold), most other i-tRNA—start codon combinations only produced modest increases in translation (Figure 2). In addition to the consistently weak ACU (i-tRNA-AGU) pair, initiation from GUU paired with i-tRNA-AAC was not detected in this strain, an occurrence unique among the strains tested.

In contrast to the other strains tested, the probiotic Nissle 1917 strain showed distinct non-canonical translation patterns. Expression of complementary i-tRNAs produced a 193-fold increase in translation initiation from the UAA start codon (i-tRNA-UUA) and a 180-fold increase from the AGG start codon (i-tRNA-CCU), representing the strongest levels of fluorescence observed in this strain. Strikingly, the fluorescence signals from these non-canonical start codons exceeded that from the canonical AUG start codon, indicating that non-canonical initiation can, in some cases, surpass native translation efficiency in this host. Previous work has demonstrated that under optimized conditions, engineered i-tRNAs capable of initiating translation from non-canonical start codons can achieve efficiencies comparable to or exceeding canonical AUG driven initiation (*17*). These improvements were shown to depend on increased availability of key translational components, including aminoacyl-tRNA synthetases, methionyl-tRNA formyltransferase, and IF2, highlighting that initiation efficiency is strongly influenced by the cellular environment. By contrast, the elevated initiation efficiencies observed here in Nissle 1917 were achieved without targeted overexpression of these factors and using distinct i-tRNA—start codon pairs, indicating that similarly high levels of non-canonical initiation can arise from different host backgrounds. In this context, the strain-dependent differences observed here may reflect variation in these and other host-specific factors, reinforcing the concept that non-canonical translation initiation is a systems-level property that must be considered in the context of the host chassis. Notably, however, these codons also exhibited ∼21-fold higher baseline expression under repressed conditions relative to other start codons (Figure S2), suggesting that the observed induction may reflect a combination of elevated basal initiation and non-canonical system activity.

Previous studies have reported that canonical GFP expression capacity follows the order K-12 > BL21(DE3) > Nissle 1917(*49*). Our data corroborate these findings and extend the comparison to additional strains, yielding the order K-12 > BL21(DE3) > Nissle 1917 > C122 > W strain > Crooks for canonical AUG-driven expression. The underlying reason for these differences is still an open question, but of the three previously investigated strains, proteomic analyses showed that ribosomal protein abundance is lowest in Nissle 1917 (*49*), which may contribute to its reduced expression capacity. In contrast to canonical translation, relatively few studies have examined the global cellular response to these non-canonical translation systems, despite the potential for these systems to introduce broader changes to cellular physiology. For example, Vincent *et al.* (*15*) reported that expression of an amber i-tRNA anticodon mutant delayed the downregulation of ribosomal proteins upon entry into the stationary phase, while Scopelliti *et al.* (*19*) observed that expression of an i-tRNA-AAC mutant led to host adaptation through upregulation of the valine biosynthetic pathway, likely to accommodate increased valine demand. Together, these findings, in combination with our observations, suggest that non-canonical translation initiation systems can significantly influence host physiology and metabolic state. In this context, such systems may provide a flexible strategy for modulating protein expression capacity in strains such as Nissle 1917 and provide a simple and effective way to boost its potential as a therapeutic modality.

Because induction ratio alone does not represent practical system useability, we next examined absolute induced expression to identify which i-tRNA—start codon pairs remain the most productive in each host background. To identify i-tRNA anticodon variants that produced the strongest expression, we performed a two-way ANOVA on induced conditions only. This analysis revealed that absolute expression remains strongly dependent on both the i-tRNA (p < 0.01) and the strain it is expressed in (p < 0.01), as well as the interaction between these two variables (p < 0.01) (Table S3).

To visualize the host-dependent trends for induced reporter fluorescence, mean fluorescence values were first calculated for each strain and i-tRNA—start codon combination. Row-wise Z-scores were then computed within each of the i-tRNA—start codon pairs across strains, allowing comparison of the relative performance of each pair across host backgrounds independent of absolute signal magnitude. This normalization highlights whether a given strain performs above or below the codon-specific average, rather than reflecting absolute differences between i-tRNA—start codon pairs. Distinct performance profiles were observed for each of these pairs, with some i-tRNAs maintaining consistently relatively high expression across strains, while others exhibited strong host-dependent shifts in activity (Figure 3A). For instance, the majority of i-tRNA—start codon pairs showed reduced relative performance in the W and Crooks strains, indicating that these hosts provide less permissive translational environments for non-canonical initiation. By contrast, the K-12 and B strains exhibited more moderate but consistent performance across i-tRNA—start codon pairs, indicating a comparatively permissive translational landscape. Conversely, the Nissle 1917 and C strains displayed greater variability in relative expression, with certain pairs showing strong initiation while others exhibit comparatively low expression. This pattern suggests that these hosts exert a stronger modulatory influence on non-canonical initiation efficiency, amplifying differences between anticodon variants.

**Figure 3.**
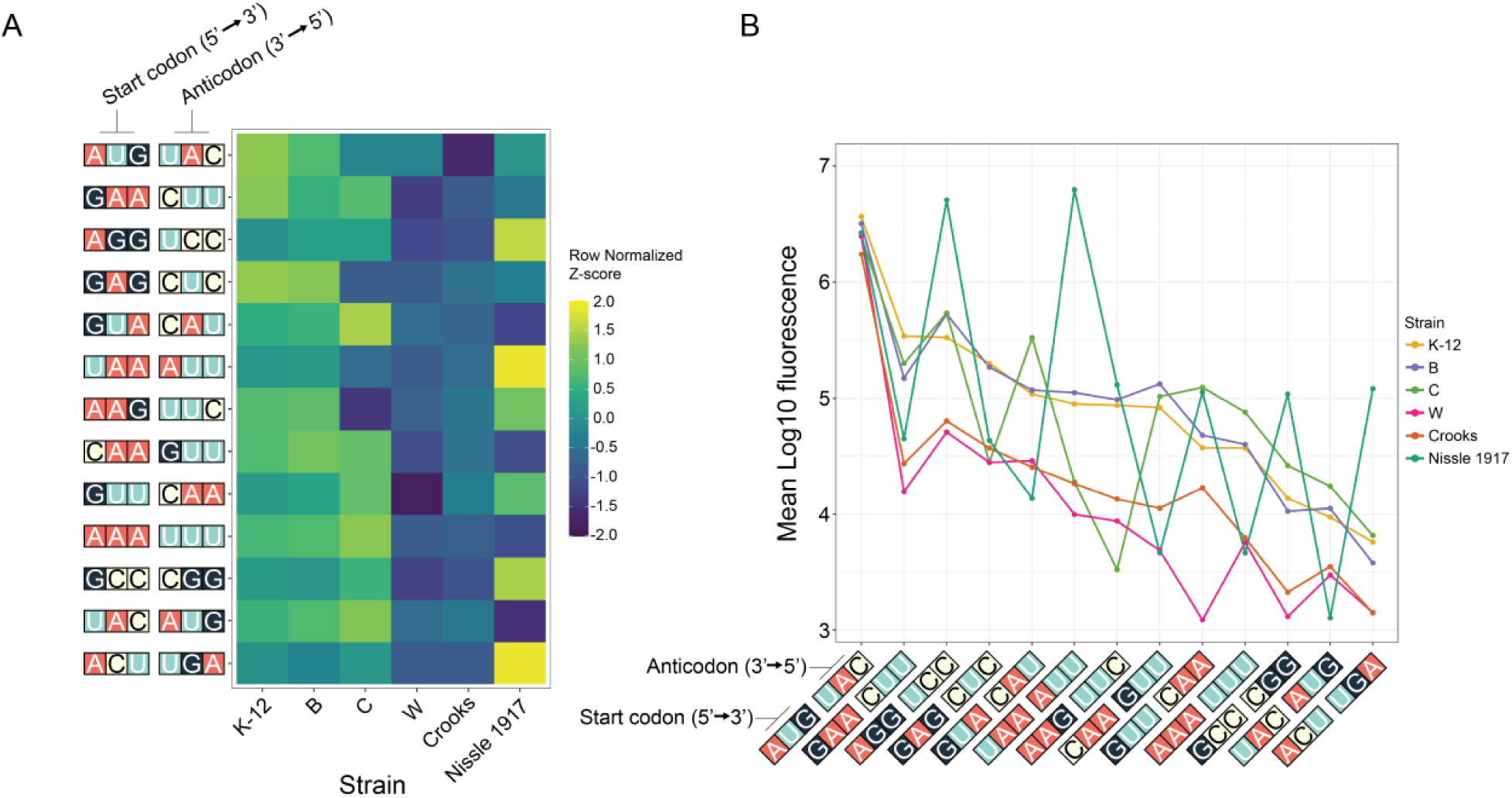
Effect of interaction between strain and mutant i-tRNAs on initiation efficiency. *E. coli* strains (K-12, B, C, W, Crooks, and Nissle 1917) expressing 12 sfGFP-i-tRNA pairs were grown in biological triplicate to mid-log phase (0.6 OD_600_) and normalized bulk fluorescence (Arb. U./OD_600_) was recorded in either repressed (2% v/v glucose) or induced (1mM IPTG) conditions. **(A)** Heatmap of row-wise Z-scores calculated from mean induced fluorescence values (Arb. U./OD_600_) for each i-tRNA—start codon pair across the six strains. Z-scores were calculated within i-tRNA—start codon pair across strains to represent relative performance independent of absolute signal magnitude where each row should be interpreted independently. **(B)** Interaction plot showing mean induced fluorescence for each i-tRNA—start codon pair across strains.

Interaction plots of mean induced fluorescence revealed pronounced changes in the rank order of mutant i-tRNA—start codon performance across strains, further illustrating the strong strain and i-tRNA—start codon pair interaction identified by two-way ANOVA (Figure 3B). The frequent crossing of lines in this plot indicates a non-parallel response pattern between strains, a hallmark of strong interaction effects. Substantial reordering of i-tRNA—start codon performance was observed depending on host background, indicating that the relative efficiency of individual variants was not conserved across strains. For example, in Nissle 1917, five of the twelve pairs ranked among the highest performing variants, while four ranked among the weakest across the dataset. This wide distribution of performance within a single host demonstrates the broad dynamic range of system behavior specific to translational context.

Across all strains, consistent trends were also observed for certain pairs. Start codons GCC, UAC, and ACU were among the poorest performing across all hosts, indicating that these i-tRNA—start codon combinations are intrinsically inefficient within the bacterial initiation framework. By contrast, the AGG start codon paired with the i-tRNA-CCU consistently produced the highest levels of expression across all strains, identifying it as a robust and broadly compatible engineered initiation pair.

Together, these analyses demonstrate that while certain anticodon variants, such as CCU, exhibit robust and portable performance, the efficiency of many others are subjected to strong host-dependent effects as a result of the translational environment. Collectively, these results demonstrate that while orthogonal translation initiation systems are broadly portable across *E. coli* lineages, their efficiency and regulatory behavior are shaped by both strain-specific translational contexts and the identity of the i-tRNA—start codon pair.

### Variation in host tRNA gene landscape influences non-canonical translation initiation efficiency

Having established strong host dependence in non-canonical initiation performance, we next asked whether comparative differences in genome content and endogenous tRNA gene repertoires might identify host features associated with strain-dependent translational performance. We first conducted a multiple whole-genome alignment using the progressiveMauve algorithm (*50*) (Figure 4A). As expected, the genomes exhibit a high degree of overall conservation, reflected by the presence of large collinear blocks representing conserved genomic segments across strains. Despite overall genomic conservation across the strains analyzed, several regions of divergence were identified that may have contributed to the observed differences in non-canonical translation initiation. Notably, Nissle 1917 contained the greatest number of strain-specific genomic regions, as indicated by gaps between conserved blocks in the multiple genome alignment. This observation is consistent with its larger genome (∼5.4 Mb) compared with the other strains analyzed here (4.6—4.9 Mb) and suggests an expanded genetic repertoire that may influence its translation capacity and regulatory landscape.

**Figure 4.**
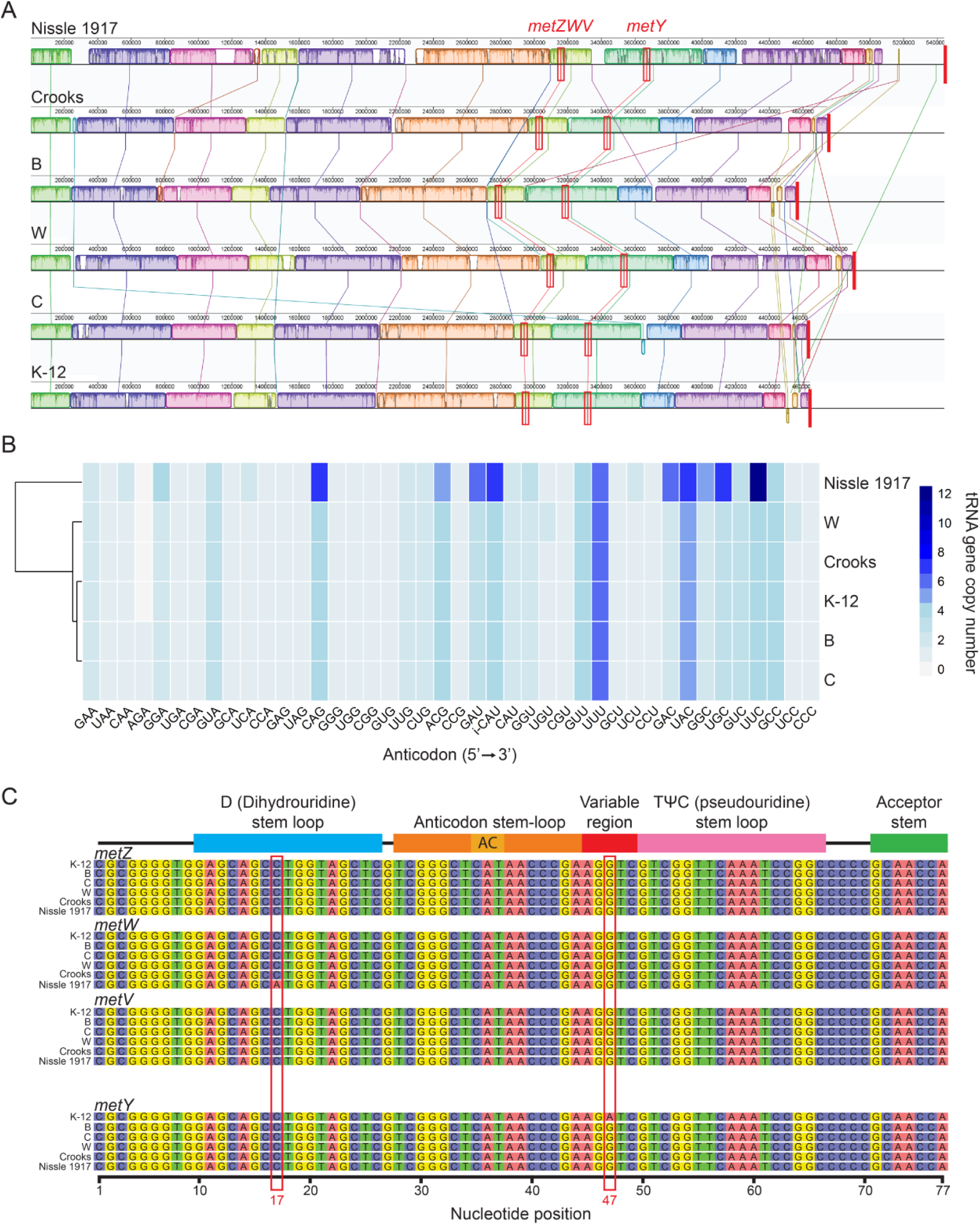
Genome and i-tRNA gene comparison across the 6 different strains of *E. coli*. **(A)** Multiple whole genome alignment of the 6 *E. coli* strains (K-12, B, C, W, Crooks, and Nissle 1917). Homologous genome regions known as locally collinear blocks (LCBS) are colored with vertical lines indicating aligned LCBs across genomes. Sequences unique to a genome are indicated by the white spaces within or between the colored LCBs. Location of the *metZWV* operon and *metY* gene across the genome are highlighted in red boxes. Red bars at the end of each genome indicate the end of each sequence. **(B)** Heatmap of tRNA gene copy numbers retrieved from the GtRNAdb across the six *E. coli* strains. i-CAT represents the native initiator tRNA. **(C)** Sequence alignment of *metZ*, *metW*, *metV*, and *metY* genes across the strains of *E. coli*. Schematic above the alignment shows the different regions with an i-tRNA sequence while axis under the alignment shows nucleotide position. Region labelled AC within the anticodon stem-loop defines location of the anticodon. Nucleotide position of sequence differences of the *metW* gene in Nissle 1917 strain and *metY* gene in K-12 MG1655 is illustrated by the red boxes.

Of particular interest, one divergent region in Nissle 1917 is located between i-tRNA gene loci, encompassing the *metZWV* operon and the *metY* gene, which encode for the native i-tRNAs. Variation within or surrounding these loci, including differences in local regulatory elements such as promoters, transcription factor binding sites, or other non-coding sequences, could plausibly influence the expression, processing, or regulation of endogenous i-tRNAs (*51*), ultimately altering the cellular tRNA pool. Because translation initiation is highly sensitive to both availability and identity of i-tRNAs, such differences may contribute to the altered initiation efficiency observed in this strain. Pan genomic analysis revealed a core genome of 3,055 genes shared across all six strains, while the accessory genome contained numerous strain-specific genes. The number of unique genes ranged from 172 in the C strain to 1,219 in Nissle 1917 (Figure S3). Functional annotations of these unique genes revealed a variety of biological functions, including type VI secretion system components, colibactin biosynthesis, and iron acquisition pathways (Supplementary File S1).

Because mutant i-tRNA-mediated initiation efficiency differed significantly across strains, we next investigated whether differences in the endogenous tRNA landscape could contribute to these host-dependent effects. Since tRNA gene copy number across a genome has been shown to correlate with tRNA abundance and forms the basis of quantitative models of translational efficiency such as the tRNA adaptation index (*52*), we assessed tRNA gene copy number as a first-order approximation of relative tRNA abundance. However, true tRNA abundance and decoding capacity can deviate from gene dosage due to growth-condition dependence, tRNA processing and maturation dynamics, and post-transcriptional modification states (*53*). Accordingly, copy number comparisons should be interpreted as indicative of potential differences in cellular tRNA pools rather than direct measurements of functional tRNA abundance. Differences in tRNA gene copy number can influence the composition and balance of the cellular tRNA pool, which in turn may affect translational dynamics and initiation fidelity. Such variation in the endogenous tRNA landscape could therefore contribute to the strain-specific differences in engineered translation efficiency observed in this study.

To compare endogenous tRNA repertoires across the strains we retrieved tRNA copy numbers from the GtRNA database (*54, 55*). Notably, Nissle 1917 contained a markedly expanded repertoire, encoding 122 tRNA genes compared with an average of 86 in the other strains analyzed (Figure 4B). In particular, Nissle 1917 encodes 12 copies of elongator tRNA-UUC, compared with only 4 copies in other *E. coli* strains. Because this elongator tRNA anticodon differs by a single nucleotide from the engineered i-tRNA-UUA used with the UAA reporter, this raises the possibility that endogenous elongator tRNAs could contribute to the unexpectedly high background initiation at this start codon (Figures 2 and S2). Prior work has shown that changes in the balance between elongator and initiator tRNAs can influence initiation fidelity (*56*), permitting non-canonical initiation events. While not directly measured in this study, this relationship is consistent with the elevated background reporter expression observed in Nissle 1917 and suggests that differences in endogenous tRNA pools may contribute to host-dependent variability in non-canonical translation performance.

Nissle 1917 also contains a greater number of i-tRNA genes compared with other strains. Native i-tRNAs have been previously reported to exhibit a degree of promiscuous initiation from many of the 64 possible start codons, including AGG (*57*). Although these prior measurements were performed in the BL21(DE3) strain, it is possible that similar promiscuous initiation events occur in Nissle 1917 and contribute to the comparatively strong expression observed from the AGG start codon in this host. It is important to note, however, that initiation and elongation dynamics are influenced by multiple factors beyond tRNA abundance, including aminoacyl-tRNA synthetase activity, ribosome availability, and cellular metabolic state (*58, 59*). Consequently, differences in tRNA gene copy numbers likely represent only one of several factors contributing to the observed host-dependent behavior of the non-canonical translation initiation system.

Looking beyond i-tRNA genomic copy number, we performed sequence alignment of the *metZWV* operon and *metY* gene across the strains in this study. This analysis revealed that these regions are nearly identical in all strains, with only single base changes from consensus in *metY* of K-12 and *metW* of Nissle 1917 (Figure 4C). In K-12, the presence of a distinct *metY*-encoded i-tRNA isoform (i-tRNA-CAU-fMet2) containing an A46 nucleotide substitution (rather than canonical G46) has been known for several decades (*60*). However, it was only recently that work has begun exploring the physiological significance of maintaining this i-tRNA isoform, suggesting that it may be involved in sustaining cellular fitness in nutrient limited conditions (*51*). Similarly, our alignment highlighted that Nissle 1917 harbors a distinct i-tRNA encoded by *metW*, containing an A17 within the dihydrouridine (D) loop, rather than the canonical C17 (Figure 4C). The functional consequences of this isoform remain unclear. However, because the D-loop interacts with the TψC-loop to stabilize and produce the characteristic L-shape of tRNAs (*61*), this substitution could plausibly influence i-tRNA structure or conformational behavior. Whether such variation affects the initiation efficiency in Nissle 1917 remains unknown and would require direct testing. However, a BLAST search and comparison with the RNAcentral database (*62*) revealed that this i-tRNA variant is not unique to Nissle 1917 but is also present in several strains of *E. coli*, *Aeromonas*, and *Shewanella* that have been implicated in disease (*63–65*). While the functional implication of this variant remains unclear, its conservation across diverse bacterial lineages suggests that, like the K-12 *metY* isoform, the A17 substitution may confer a selective advantage under specific ecological contexts.

### Mutant i-tRNAs impose minimal fitness effects across E. coli hosts

Because practical deployment of non-canonical translation systems requires not only successful expression but also host tolerance, we next evaluated the fitness consequences of mutant i-tRNA expression across strains. To assess this, we quantified changes in growth rate and maximum optical density by comparing induced and repressed conditions for each mutant i-tRNA (Figure 5). Overall, expression of mutant i-tRNAs was well tolerated, with most strain and i-tRNA—start codon combinations exhibiting no significant fitness defects (Figure S4—S16). However, a two-way ANOVA revealed significant effects of strain (p < 0.001), i-tRNA (p < 0.01), and their interaction (p < 0.01) on growth rate (Table S4), indicating that mutant i-tRNAs can exert measurable and significant impacts on cellular fitness, but that the effects are highly dependent on host background and specific i-tRNA—start codon combinations.

**Figure 5.**
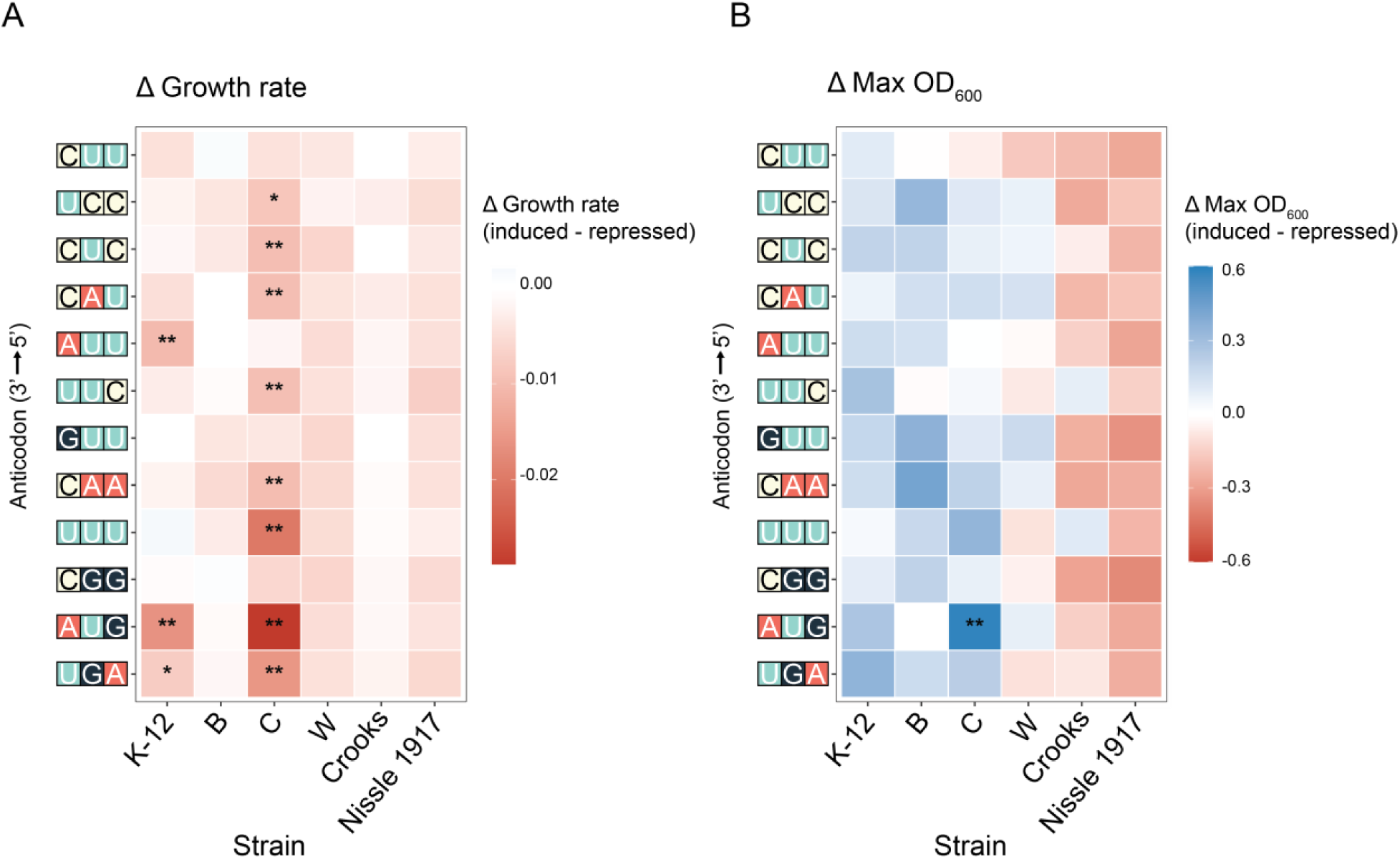
Strain dependent effects of mutant i-tRNAs on cellular fitness. Changes in growth rate (Δ growth rate) and maximum optical density (Δ MaxOD) were determined by comparing induced (1mM IPTG) and repressed (2% v/v glucose) conditions for each of the six *E. coli* strains (K-12, B, C, W, Crooks, and Nissle 1917) expressing 12 mutant i-tRNA variants. Cells were grown in biological triplicate to mid-log phase (0.6 OD_600_) and growth parameters were calculated from normalized growth curves. **(A)** Heatmap of Δ growth rate values and **(B)** Δ MaxOD values. Statistical analysis was performed using two-way ANOVA followed by Tukey’s post hoc testing where * denotes p < 0.05 and **denotes p < 0.01. Color scales represent the magnitude of Δ values, with red indicating reduced fitness and blue indicating increased fitness.

Consistent with this, reductions in growth rate due to i-tRNA variant expression were strongly strain dependent (Figure 5A). The C strain was the most sensitive host, with significant decreases observed in 8 of 12 i-tRNAs, whereas K-12 only had growth defects in 3 of 12 combinations. By contrast, Crooks, W, B, and Nissle 1917 displayed minimal sensitivity to mutant i-tRNA expression.

In contrast to growth rate, maximum culture optical density was largely unaffected by mutant i-tRNA expression. Two-way ANOVA revealed a significant effect of strain (p < 0.01), but no significant contribution from i-tRNA (p = 0.48) or the interaction between strain and i-tRNA (p = 0.80) (Table S5). This decoupling of growth dynamics and final optical density suggests that while non-canonical initiation perturbs growth rate, it does not significantly affect population size over longer timescales, potentially buffering their impact on overall population fitness. These findings support the notion that orthogonal translation initiation systems can be deployed with minimal impact on host viability in monoculture, despite measurable perturbations to translational performance.

One possibility for the observed strain-dependent fitness effects is that mutant i-tRNAs must compete with endogenous elongator tRNAs for aminoacyl-tRNA synthetase (aaRS) binding and charging. Anticodon mutated i-tRNAs can exhibit altered identity determinants and may be recognized by non-cognate aaRSs, resulting in charging with amino acids other than methionine, as has been previously demonstrated for several i-tRNA mutants (*17, 19*). Such competition could disrupt the balance of the cellular tRNA pool, increasing the proportion of uncharged endogenous tRNAs and promoting ribosomal stalling, ultimately impairing translational efficiency. The strain-specific nature of the growth defects may therefore reflect intrinsic metabolic differences and the varying capacity of each strain to aminoacylate and utilize these heterologous tRNAs (*46, 66, 67*), influencing their ability to accommodate the additional translational burden. Additionally, variations in tRNA modification profiles, aminoacylation efficiency, and ribosome binding dynamics may also affect the stability and function of mutant i-tRNAs within each strain, shaping their impact on both protein expression and cellular fitness (*68, 69*). Together, these factors may underlie the observed divergence in both non-canonical initiation efficiency and fitness effects across host backgrounds. Although these mechanisms remain to be tested directly, the data indicate that host tolerance to engineered initiation systems is not uniform and likely depends on multiple layers of strain-specific physiology.

Further studies integrating proteomic and RNA-based analyses will be important for further elucidating the mechanisms underlying these observations (*19*). In particular, examining ribosome occupancy and stalling, global tRNA aminoacylation levels, and ribosome assembly dynamics could provide insight into how mutated i-tRNAs interact with host translational machinery. Characterizing mutant i-tRNA modification profiles, aminoacylation efficiency, and charged amino acid identity will also be essential for defining the determinants of non-canonical translation efficiency across different strains. Such insights will help guide the development of improved orthogonal translation systems with enhanced performance and minimal impact on host physiology.

These combined analyses of inducibility, absolute expression, and fitness effects provide a framework for application-specific selection of host strains for orthogonal translation system deployment (Table 1). For general use, the K-12 and B strains represent the most versatile hosts, supporting robust non-canonical translation across multiple i-tRNA—start codon pairs. Within these, K-12 exhibits a broad dynamic range of expression, enabling multiplexed systems with variable output. In contrast to the K-12 strain, the B strain provides a more balanced expression profile, with several i-tRNA—start codon pairs producing similar levels of expression, making it advantageous for applications requiring coordinated or uniform protein production.

**Table 1.** Application-guided selection of host strains and non-canonical initiation pairs for optimized system performance.

| Use case | Host strain/s | Representative start codon—i-tRNA pairs | Rationale | Considerations |
| --- | --- | --- | --- | --- |
| **General purpose use across multiple non-canonical pairs** | K-12, B | GAA—i-tRNA-UUC,<br>AGG—i-tRNA-CCU,<br>GAG—i-tRNA-CUC,<br>GUA—i-tRNA-UAC,<br>UAA—i-tRNA-UUA, | Supports robust non-canonical translation across multiple i-tRNA—start codon pairs. | Elevated basal expression may reduce regulatory tightness. |
| **Multiplexed expression with variable output** | K-12 | GAA—i-tRNA-UUC,<br>AAG—i-tRNA-CUU,<br>GCC—i-tRNA-GGC | Wide dynamic range enables differential expression across pairs. | i-tRNA-GUA, i-tRNA-AGU, and i-tRNA-UUA introduced growth defects. |
| **Multiplexed expression with uniform output** | B | GAA—i-tRNA-UUC,<br>GAG—i-tRNA-CUC,<br>GUA—i-tRNA-UAC,<br>UAA—i-tRNA-UUA,<br>AAG—i-tRNA-CUU,<br>CAA—i-tRNA-UUG | Multiple non-canonical pairs produce consistent and similar expression levels. | Mild background expression. |
| **Tight regulation with low background expression** | Crooks | GAA—i-tRNA-UUC,<br>GAG—i-tRNA-CUC,<br>GUA—i-tRNA-UAC,<br>UAA—i-tRNA-UUA,<br>AAG—i-tRNA-CUU | Minimal basal expression and controlled induction. | Lower absolute expression overall compared to other strains. |
| **Maximum protein output** | Nissle 1917 | AGG—i-tRNA-CCU,<br>UAA—i-tRNA-UUA | Selected non-canonical pairs exceed canonical AUG-driven expression. | Elevated baseline expression. |

In scenarios where tight regulation and minimal background expression are prioritized over maximal output, the Crooks strain offers a favorable host, exhibiting low basal fluorescence and controlled activation of non-canonical translation. Conversely, when maximizing absolute protein output is the primary objective, Nissle 1917 emerges as a suitable host. In this strain, non-canonical start codon UAA (i-tRNA-UUA) and AGG (i-tRNA-CCU), produced expression levels exceeding those observed from the canonical AUG start codon, highlighting its potential for high yield protein production. The probiotic uses of Nissle 1917 further expands its applicability in therapeutic and clinical contexts (*70*).

Together, these analyses show that non-canonical translation initiation cannot be evaluated solely on the basis of system output in an individual host. Instead, non-canonical translation initiation system deployment reflects a multifactorial trade-off between inducibility, absolute output, regulatory tightness, and physiological tolerance, each of which is shaped by host background. These results therefore support a host-aware framework for selecting appropriate strain—system combinations according to the requirements of a given synthetic biology application.

## Conclusions

In this study, we systematically evaluated the portability of a non-canonical translation initiation system across six commonly used *E. coli* GRAS strains. While non-canonical initiation was broadly functional, its performance varied substantially depending on both the host background and the i-tRNA—start codon pair. Inducibility, absolute expression output, and host fitness effects were not uniformly consistent across strains, demonstrating that these translation systems cannot be assumed to function equivalently across different chassis organisms.

These findings establish host background as a key design variable for non-canonical translation system deployment. Rather than relying on single-strain benchmarking, effective implementation requires consideration of multiple performance parameters, including expression strength, regulatory tightness, and physiological tolerance. The framework presented here enables application-specific selection of host—system combinations, supporting use cases ranging from multiplex expression to high-yield protein production.

More broadly, this work highlights that translational performance and host tolerance are not inherently coupled and that predicting orthogonal system behavior will require a deeper understanding of host-specific translational physiology. While comparative genomic and tRNA landscape analyses identify candidate contributors to these differences, further studies integrating tRNA abundance, aminoacylation state, and global translation dynamics will be necessary to define the underlying mechanisms. Together, these results underscore the importance of host-aware engineering strategies for achieving predictable and scalable control of translation across chassis organisms used in synthetic biology.

## Methods

### Bacterial strain selection and growth conditions

All *E. coli* strains used are commonly recognized as generally safe for research strains with designated risk group 1. In total, six strains were selected for further study including MG1655 (K-12, ATCC #47076), B (ATCC #BAA-1025), C (NCTC122), Crooks (ATCC #8739), W (ATCC #9637), and Nissle 1917(*33, 71–75*). All strains within this study were grown in lysogeny broth Miller (LB^M^)(10 g/L Peptone, 10 g/L NaCl, 5 g/L yeast extract and Milli-Q^®^ Ultrapure water) or streaked and grown on LB^M^ agar (10 g/L Peptone, 10 g/L NaCl, 5 g/L yeast extract and Milli-Q^®^ Ultrapure water, 15 g/L agar), supplemented with appropriate antibiotics. Spectinomycin (50 µg/mL) and carbenicillin (100 µg/mL) were used concurrently to maintain pULTRA-tac-*metY* and pQE-T5-*sfGFP* plasmids, respectively. Overnight cultures were inoculated in 1 mL of LB^M^ with appropriate antibiotics and grown overnight in an InforsMT Multitron pro-orbital shaker-incubator for 16 hours at 37°C with shaking (250 rpm). Cultures were then back diluted to an OD_600_ of 0.1 in fresh LB^M^ and grown under experimental conditions as described below for each assay.

### Plasmid construction

To create the pULTRA-*tac*-*metY* plasmids, the pULTRA-CNF vector, which was a gift from Peter Schultz (Addgene plasmid # 48215) (*76*), was linearized by inverse PCR and ligated with synthetic DNA (Integrated DNA technologies) containing anticodon mutant *metY* genes using Gibson assembly (NEBuilder HiFi DNA Assembly kit (NEB#E2621)). Anticodon mutants included TTC, CCT, CTC, TAC, TTA, CTT, TTG, AAC, TTT, GGC, GTA, or AGT (Table S6). The 12 pQE60-*T5-sfGFP* reporter plasmids used in this study were created through site-directed mutagenesis using the original pQE60-*T5-sfGFP(AUG)* plasmid obtained from a previous study (*47*) as a template (Table S6). The original template was linearized using Q5^®^ Hot Start High-Fidelity 2X Master Mix (NEB #M0494) and a suite of non-overlapping primers designed to introduce the desired mutations to the start codons of the sfGFP reporter protein (Table S7). The resulting linear PCR products were subjected to DpnI (NEB #R0176S) digestion to remove the original template then re-circularized using T4 DNA ligase (NEB #M0202) to generate pQE60*-T5-sfGFP* plasmids with start codons AAA, AAG, ACU, AGG, CAA, GAA, GAG, GCC, GUA, GUU, UAA, UAC. The resulting pQE60-*T5-sfGFP* plasmids were then sequence verified using pQE60_SEQ_FWD and pQE60_SEQ_REV primers (Table S7). All plasmids harboring complementary *metY* anticodons and sfGFP start codons were co-transformed into respective bacterial strains using the Mix & Go Transformation Kit^®^ (Zymo Research cat #T3002) as per manufacturer instructions.

### Fluorescence measurements

Bacterial starter cultures containing the variants of the pULTRA-*tac-metY* plasmid along with complementary versions of the pQE60-*T5-sfGFP* were established as described above. These cultures were then back diluted 1:100 in 800 µL of fresh LB^M^ with appropriate amounts of carbenicillin and spectinomycin then grown for 2 hours at 37 °C with shaking. Biological triplicates from each culture were then split into paired conditions: (1) test condition induced with 1 mM IPTG, and (2) control condition repressed with 2% (v/v) glucose; and growth for a further 4 hours at 37 °C. Following induction, 300 µL of bacterial culture from each condition was transferred into clear 96-well plates (Sigma, cat#CLS3610) and centrifuged in a swinging bucket rotor at 2,240 g for 13 minutes at 4°C. Subsequently, the supernatant was removed, and the pelleted cells were resuspended in 200 µL of phosphate-buffered saline (PBS) and incubated overnight at 4°C to allow for sfGFP maturation. The next day, the resuspended pellet was transferred into a black clear bottom 96-well plates (Sigma, cat#CL3603) in preparation for fluorescence measurements. Absorbance (OD_600_) and bulk fluorescence for sfGFP expression was measured using a BioTek Synergy H1 plate reader (excitation 488 nm, emission 530 nm). Results were recorded with a variable gain setting to accommodate the high dynamic range of sfGFP fluorescence. Fluorescence values were normalized to OD_600_ to account for differences in cell density.

### Fitness Analysis

Conducted as previously described by (*15*), (*47*) with modifications. Bacterial starter cultures harbouring pULTRA-*tac-metY* variants were established as described above. These strains were then back diluted 1:100 into 1 mL of fresh LB^M^ with antibiotics and grown at 37°C with shaking for 4 hours in 2 mL 96 DeepWell^®^ growth plates (Merck cat# Z717266). Subsequently, cells were back diluted to 0.1 OD_600_ with a final volume of 200 µL of LB^M^ with appropriate antibiotics in a flat bottom 96-well plate and sealed with a gas-permeable transparent seal (Sigma cat# Z380059). Biological replicates were then split into two paired growth conditions: (1) an induced condition with 1 mM IPTG, and (2) control condition repressed with 2% (v/v) glucose. Both conditions were grown for 18 hours at 37 °C in a BioTek Synergy H1 plate reader, with absorbance at OD_600_ readings occurring every 5 minutes, and continuous shaking.

### Multiple sequence alignment and phylogenetic tree building

A phylogenetic tree of 18 different *E. coli* strains and 2 *Escherichia fergusonii* strains was constructed using the tree tool in Geneious Prime 2024.0.5 (https://www.geneious.com). Whole genome sequences were downloaded from NCBI genome database. Sequences used include: *Escherichia fergusonii* ATCC 35469 (NC_011740.1), *Escherichia fergusonii* FDAARGOS_1499 (NZ_CP083638.1), SMS-3-5 (NC_010498.1), UMN026 (NC_011751.1), 55989 (NZ_CP028304.1), O127:H6 str. E2348/69 (NC_011601.1), O6:K15:H31 536 (NC_008253.1), UTI89 (NC_007946.1), CFT073 (NC_004431.1), Nissle 1917 (NZ_CP007799.1), SE11 (NC_011415.1), W ATCC9637 (NC_017664.1), Crooks ATCC8739 (NC_010468.1), NCTC122 (LT906474.1), K-12 substr. MG1655 (NC_000913.3), K-12 substr. W3110 (NC_007779.1), ETEC H10407 (NC_017633.1), BL21(DE3) (NC_012971.2), 139 (NZ_CP028620.1), O157:H7 str. Sakai (NC_002695.2). MLST analysis was conducted using MLST2.0.9 on the Center of Genomic Epidemiology platform (https://cge.food.dtu.dk/services/MLST/) (*77*) with the MLST configuration set to Escherichia coli#1 which included *adk*, *recA*, *purA*, *fumC*, *gyrB*, *mdh*, *icd* genes. MLST allele sequences were then downloaded and imported into Geneious Prime 2024.0.5 (https://www.geneious.com). The sequences were concatenated and a multiple sequence alignment using the MUSCLE v5.1 (*78*) was performed. The alignment was used to create a phylogenetic tree with the Geneious Tree Builder tool using the neighbor-joining tree building method employing the Tamura-Nei genetic distance model. A Bootstrap method was used to resample the tree with a total number of 1000 replicates.

### Genomic analysis and comparison

To carry out a multiple sequence alignment of complete genomes to compare the strains selected in this study, the NCBI sequences of Nissle 1917 (NZ_CP007799.1), W ATCC9637 (NC_017664.1), Crooks ATCC8739 (NC_010468.1), NCTC122 (LT906474.1), K-12 substr. MG1655 (NC_000913.3), and BL21(DE3) (NC_012971.2) that were imported into Geneious Prime 2024.0.5 (https://www.geneious.com) were aligned and visualized using the Mauve plug-in v1.1.3 using the progressiveMauve algorithm with default parameters (*50, 79*). Comparison of annotated genes were carried out by downloading the annotation features (GFF) and genome sequence (FASTA) files for each of the six strains from the NCBI genome database. The two files were manually concatenated to generate a GFF3 file which was then analyzed using Roary v3.13.0 (*80*) using a 95% minimum identity cutoff. The output file “gene_presence_absence.csv” (File S1) was converted into a binary matrix using Microsoft Excel and visualized using the R package UpSetR (*81*). Sequences for the *metZWV* operon and *metY* gene were extracted from whole genome sequences in Geneious Prime 2024.0.5 (https://www.geneious.com) and aligned using MUSCLE v5.1 algorithm (*78*). Comparison of tRNA gene sets was carried out using the Genomic tRNA database (GtRNAdb) release 21 (http://gtrnadb.ucsc.edu/) (*55, 82*).

### Data analyses

All statistical analyses were performed in R (v2025.05.1). For fluorescence-based reporter assays, two complementary analytical approaches were used to assess system performance. To evaluate inducibility, fluorescence values were first normalized to OD_600_ to account for differences in cell density. Log2 fold change values were then calculated for each biological replicate (n=3) using the paired induced and repressed measurements obtained from the same starter cultures. Log2-transformed fold change values were analyzed using a two-way ANOVA with strain and i-tRNA—start codon pair as fixed factors. To assess absolute non-canonical system output independent of basal expression, OD_600_ normalized fluorescence values from induced conditions alone were analyzed separately using a second two-way ANOVA. For both analyses, post hoc pairwise comparisons were performed using Tukey’s (HSD) test. Estimated marginal means were calculated using the *emmeans* package (v2.0.2), and compact letter displays were generated using the *multcompView* package (v0.1-11) to indicate statistically significant groupings. Growth rate (μr) and maximum cell optical density (maxOD_600_) were analyzed using the GrowthCurver R package (*83*). Difference in growth parameters between induced and repressed conditions (ΔGrowth rate and ΔmaxOD) were calculated for each biological replicate and analyzed using a two-way ANOVA followed by Tukey’s HSD post hoc testing.

## Abbreviations

i-tRNAs: Initiator tRNAs
maxOD: Maximum OD600
IFs: Initiation factors
SD: Shine-Dalgarno
fMet: Formyl-methionine
30S PIC: 30S pre-initiation complex
30S IC: 30S initiation complex
aaRSs: Aminoacyl tRNA synthases

## Author Contributions

**Dominic Scopelliti (DS):** Conceptualization, Experimentation, Formal Analysis, Investigation, Methodology, Visualization, Writing - Original Draft, Writing - Review.

**Andras Hutvagner (AH):** Conceptualization, Experimentation, Formal Analysis, Investigation, Methodology, Visualization, Writing – Original Draft, Funding Acquisition.

**Paul R. Jaschke (PRJ):** Conceptualization, Writing - Review & Editing, Supervision, Project administration, Funding acquisition.

## Acknowledgements

We would like to thank Ariel Hecht, Sasha Tetu and Russel Vincent for the R code required for data analysis and the production of several figures. Throughout the duration of creating and writing this manuscript DS was supported by a Macquarie Research Excellence PhD Scholarship, AH was supported by a Macquarie Research Excellence PhD Scholarship and CSIRO SynBio FSP Top-Up scholarship, and PRJ was supported by the Molecular Sciences Department, Faculty of Science & Engineering, and the Deputy Vice-Chancellor Research (DVCR) of Macquarie University.

