## Supplemental for "Host background shapes the portability of a non-canonical translation initiation system across *Escherichia coli* strains"

Contents

**Supporting Figures**

Figure S1	Strain-dependent differences in basal sfGFP expression under repressed conditions.

Figure S2	Strain-dependent differences in basal sfGFP expression across anticodon—start codon pairs under repressed conditions.

Figure S3	Pan-genome comparison of the six *E. coli* strains.

Figure S4	Fitness analysis of top performing i-tRNA mutants expressed within a suite of six laboratory strains.

Figures S5-S16	Growth curves and fitness analysis of each i-tRNA mutant expressed within each strain analyzed within this study.

**Supporting Tables**

Table S1	Two-way ANOVA of log_2_ fold-change in fluorescence

Table S2	Two-way ANOVA of repressed-condition fluorescence

Table S3	Two-way ANOVA of induced-condition fluorescence

Table S4	Two-way ANOVA of growth rate

Table S5	Two-way ANOVA of maximum optical density

Table S6	Gene sequences

Table S7	Oligos used

**Supporting Files**

Supporting File S1	Gene_presence_absence

Supporting File S2 	Bulk fluorescence data

Supporting File S3 	Fitness assay data

Supporting File S4	Annotated plasmid sequence: pULTRA-tac*-metY*(NNN)

Supporting File S5	Annotated plasmid sequence: pQE60-T5-(NNN)*sfGFP*

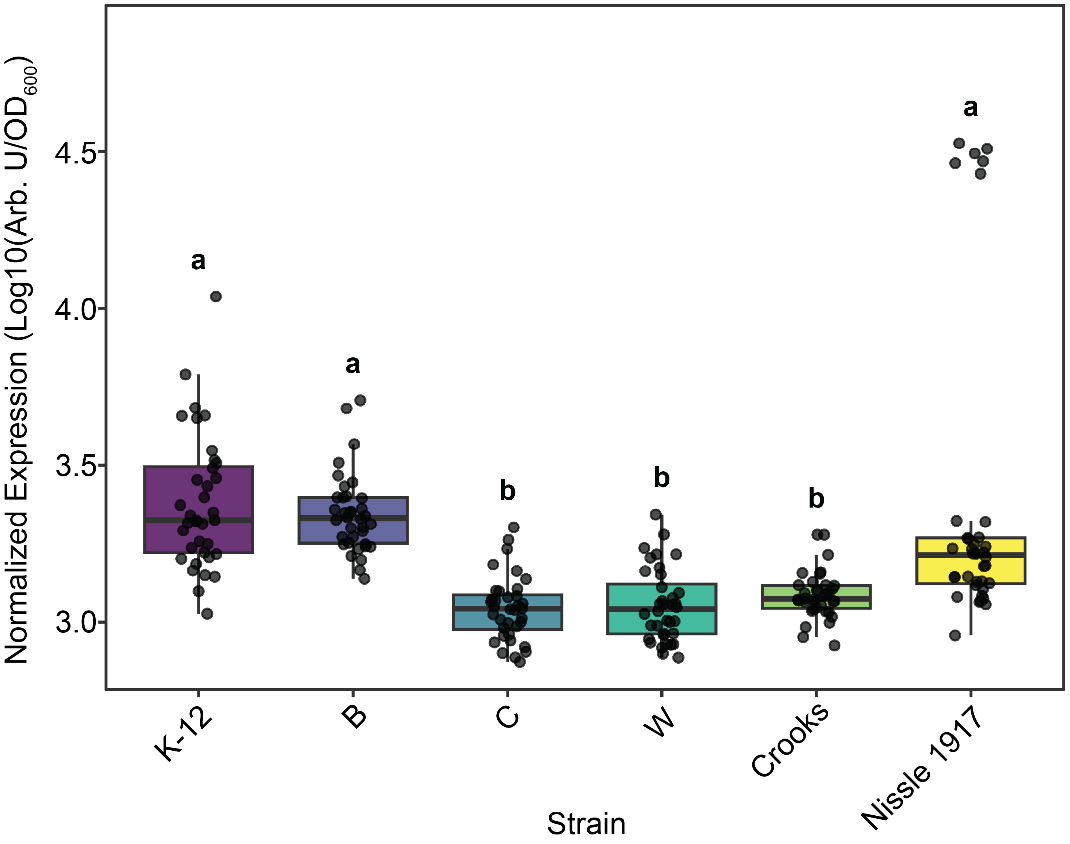

**Figure S1. Strain-dependent differences in basal sfGFP expression under repressed conditions.** Normalized fluorescence (Arb. U./OD_600_) was measured for each of the six *E. coli* strains (K-12, B, C, W, Crooks, and Nissle 1917) harboring the orthogonal translation initiation system under repressed conditions (2% v/v glucose). Cells were grown to mid-log phase (0.6 OD_600_), and fluorescence values represent background expression in the absence of IPTG induction. Individual data points represent biological replicates pooled across anticodon—start codon pairs within each strain. Boxplots showing median and interquartile range. Statistical analyses were performed using a two-way ANOVA followed by Tukey’s post hoc test. Different letters above each group indicate statistically indistinguishable strains (p < 0.05).

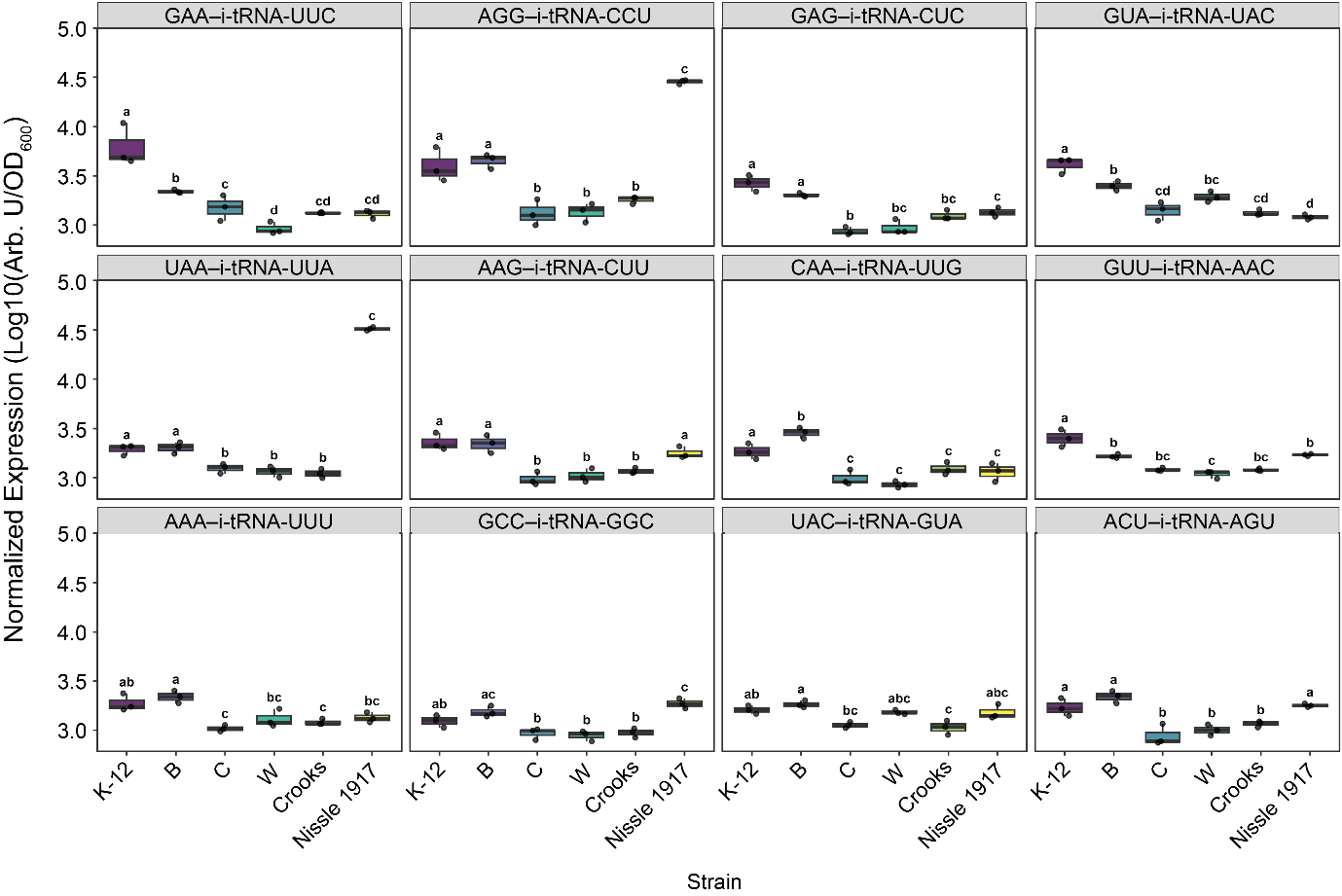

**Figure S2. Strain-dependent differences in basal sfGFP expression across anticodon—start codon pairs under repressed conditions.** Normalized fluorescence (Arb. U./OD_600_) was measured for each of the six *E. coli* strains (K-12, B, C, W, Crooks, and Nissle 1917) harboring the orthogonal translation initiation system under repressed conditions (2% v/v glucose). Cells were grown to mid-log phase (0.6 OD_600_), and fluorescence values represent background expression in the absence of IPTG induction. Each panel corresponds to a specific anticodon—start codon pair. Individual data points represent biological replicates for each strain. Boxplots display the median and interquartile range. Statistical analysis was performed using a two-way ANOVA followed by Tukey’s post hoc test. Different letters above each group indicate statistically indistinguishable strains (p < 0.05).

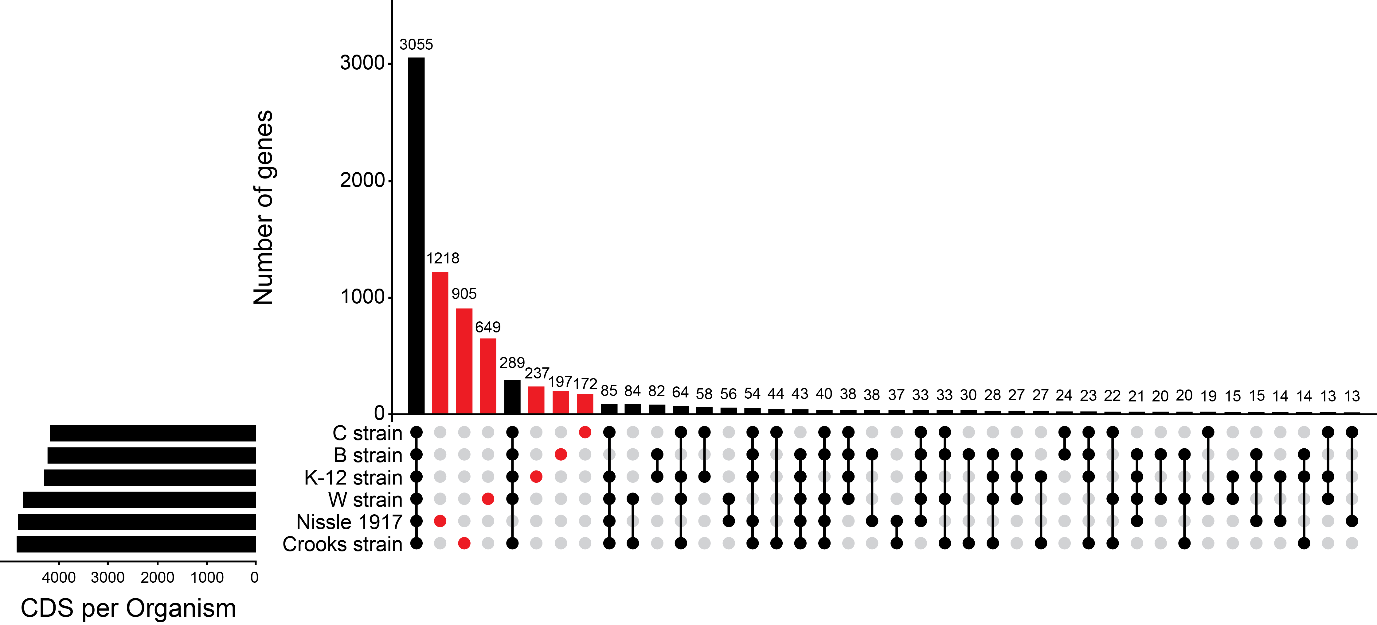

**Figure S3. Pan-genome comparison of the six *E coli* strains.** Upset plot generated using Roary v3.13.0 analysis to highlight unique genes (red) and shared genes (black) across strains. Present and absent genes were extracted from NCBI genome assemblies of Nissle 1917 (NZ_CP007799.1), W ATCC9637 (NC_017664.1), Crooks ATCC8739 (NC_010468.1), NCTC122 (LT906474.1), K-12 substr. MG1655 (NC_000913.3), and BL21(DE3) (NC_012971.2).

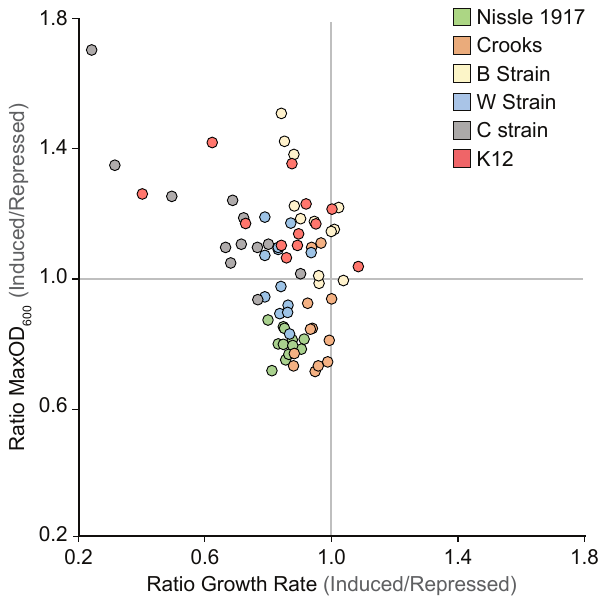
**Figure S4. Fitness analysis of six laboratory strains singly expressing one of 12 mutant i-tRNAs.** Ratios of growth rate and maximum cell density were determined for strains harboring pULTRA-*tac-metY* variants. All strains were grown in LB in the presence of spectinomycin, induced with 1 mM IPTG, or repressed with 2% v/v glucose (n=3).

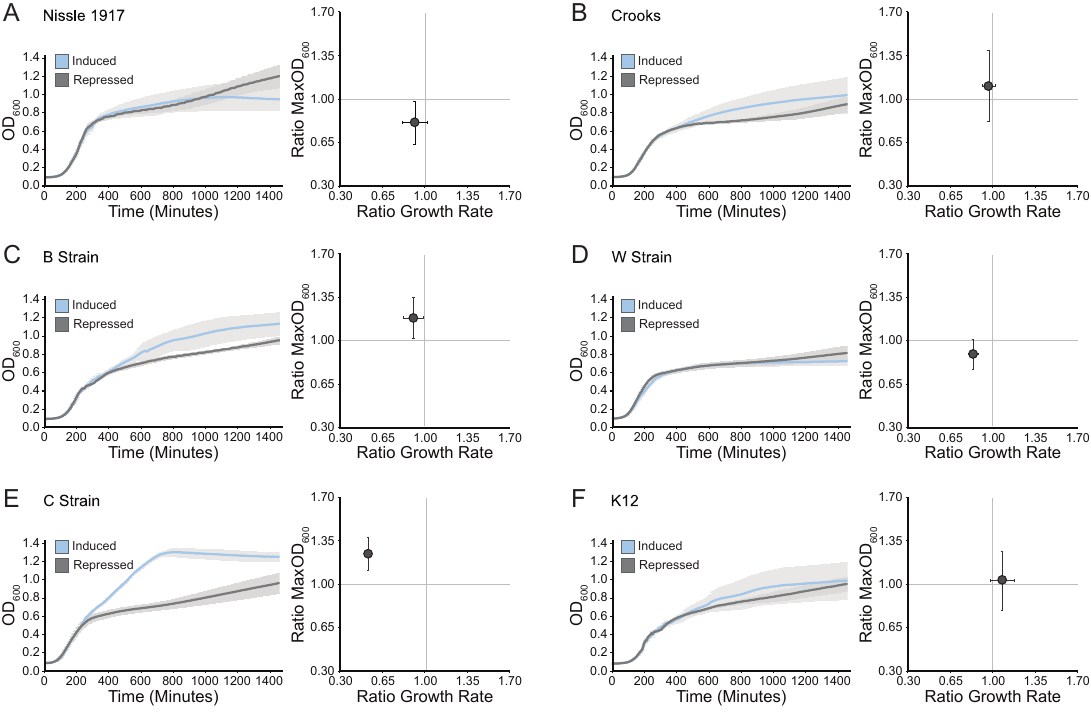
**Figure S5. Fitness analysis of six common laboratory strains expressing the mutant initiator tRNA-UUU.** Growth rate and maximum cell density were determined for strains grown in LB (spec) and induced with 1 mM IPTG, or repressed with 2% v/v glucose (n=3). **(Left)** Growth of cells carrying i-tRNA-UUU in an induced (blue) versus repressed (grey) condition. Line is the average value (n=3) and shading indicates bounds of one standard deviation. **(Right)** Growth rate and maximum cell density of cells expressing i-tRNA-UUU. Point represents average value and error bars represent one standard deviation (n=3).

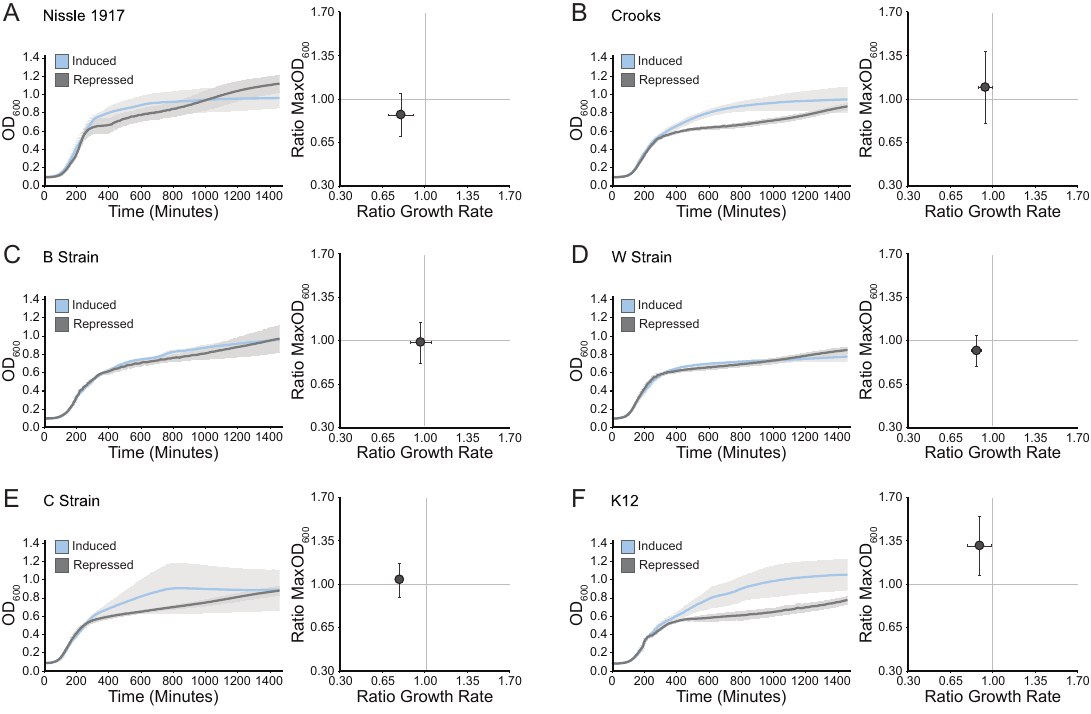
**Figure S6. Fitness analysis of six common laboratory strains expressing the mutant initiator tRNA-CUU.** Growth rate and maximum cell density were determined for strains grown in LB (spec) and induced with 1 mM IPTG, or repressed with 2% v/v glucose (n=3). **(Left)** Growth of cells carrying i-tRNA-CUU in an induced (blue) versus repressed (grey) condition. Line is the average value (n=3) and shading indicates bounds of one standard deviation. **(Right)** Growth rate and maximum cell density of cells expressing i-tRNA-CUU. Point represents average value and error bars represent one standard deviation (n=3).

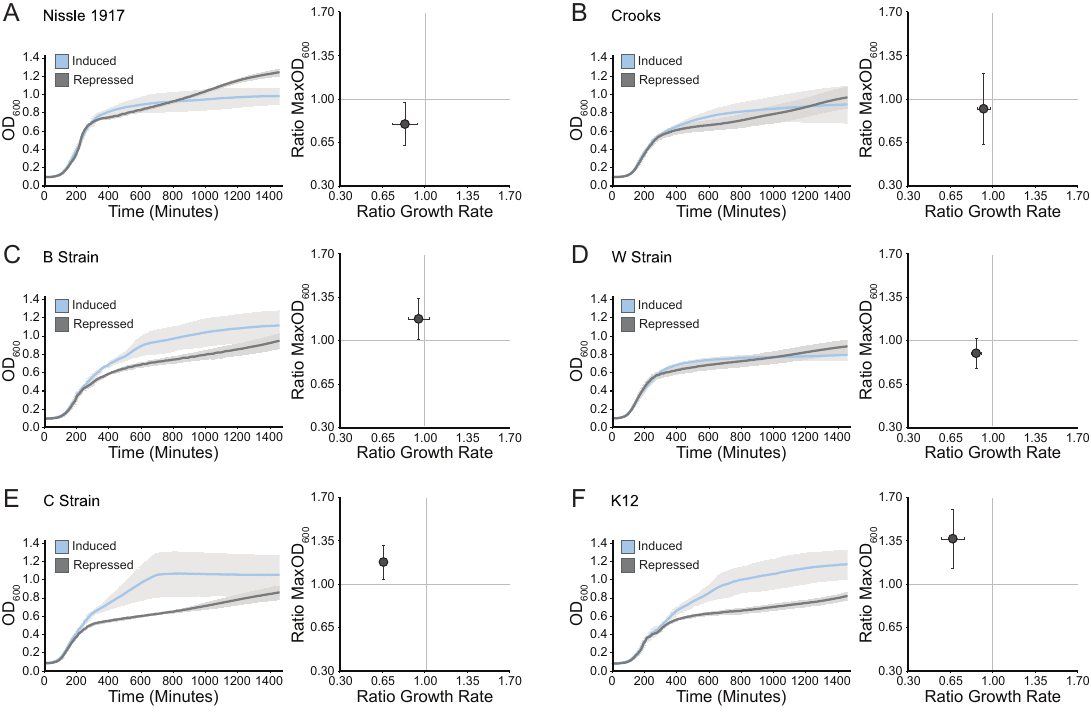
**Figure S7. Fitness analysis of six common laboratory strains expressing the mutant initiator tRNA-AGU.** Growth rate and maximum cell density were determined for strains grown in LB (spec) and induced with 1 mM IPTG, or repressed with 2% v/v glucose (n=3). **(Left)** Growth of cells carrying i-tRNA-AGU in an induced (blue) versus repressed (grey) condition. Line is the average value (n=3) and shading indicates bounds of one standard deviation. **(Right)** Growth rate and maximum cell density of cells expressing i-tRNA-AGU. Point represents average value and error bars represent one standard deviation (n=3).

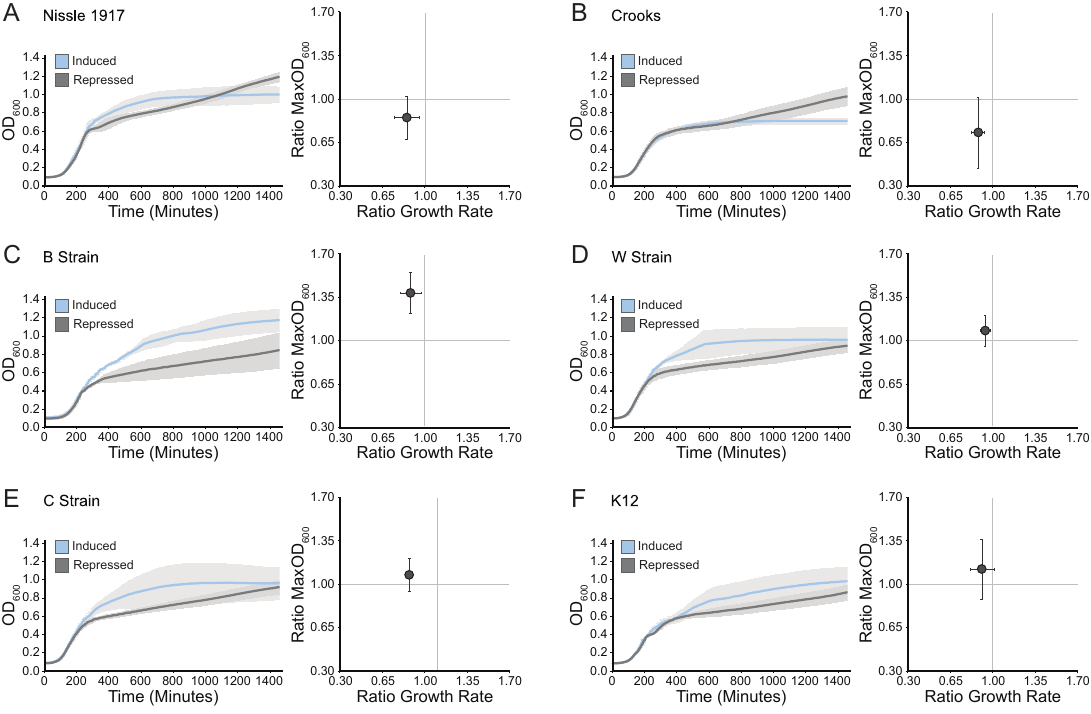

**Figure S8. Fitness analysis of six common laboratory strains expressing the mutant initiator tRNA-CCU.** Growth rate and maximum cell density were determined for strains grown in LB (spec) and induced with 1 mM IPTG, or repressed with 2% v/v glucose (=3). **(Left)** Growth of cells carrying i-tRNA-CCU in an induced (blue) versus repressed (grey) condition. Line is the average value (n=3) and shading indicates bounds of one standard deviation. **(Right)** Growth rate and maximum cell density of cells expressing i-tRNA-CCU. Point represents average value n and error bars represent one standard deviation (n=3).

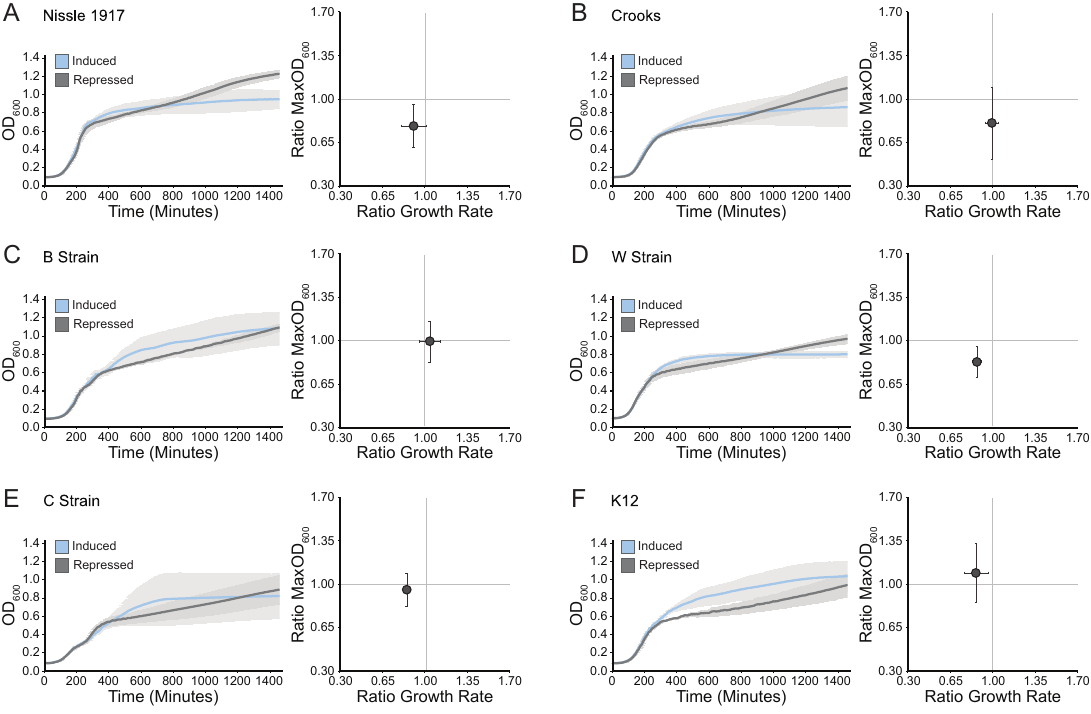
**Figure S9. Fitness analysis of six common laboratory strains expressing the mutant initiator tRNA-UUG.** Growth rate and maximum cell density were determined for strains grown in LB (spec) and induced with 1 mM IPTG, or repressed with 2% v/v glucose (n=3). **(Left)** Growth of cells carrying i-tRNA-UUG in an induced (blue) versus repressed (grey) condition. Line is the average value (n=3) and shading indicates bounds of one standard deviation. **(Right)** Growth rate and maximum cell density of cells expressing i-tRNA-UUG. Point represents average value n and error bars represent one standard deviation (n=3).

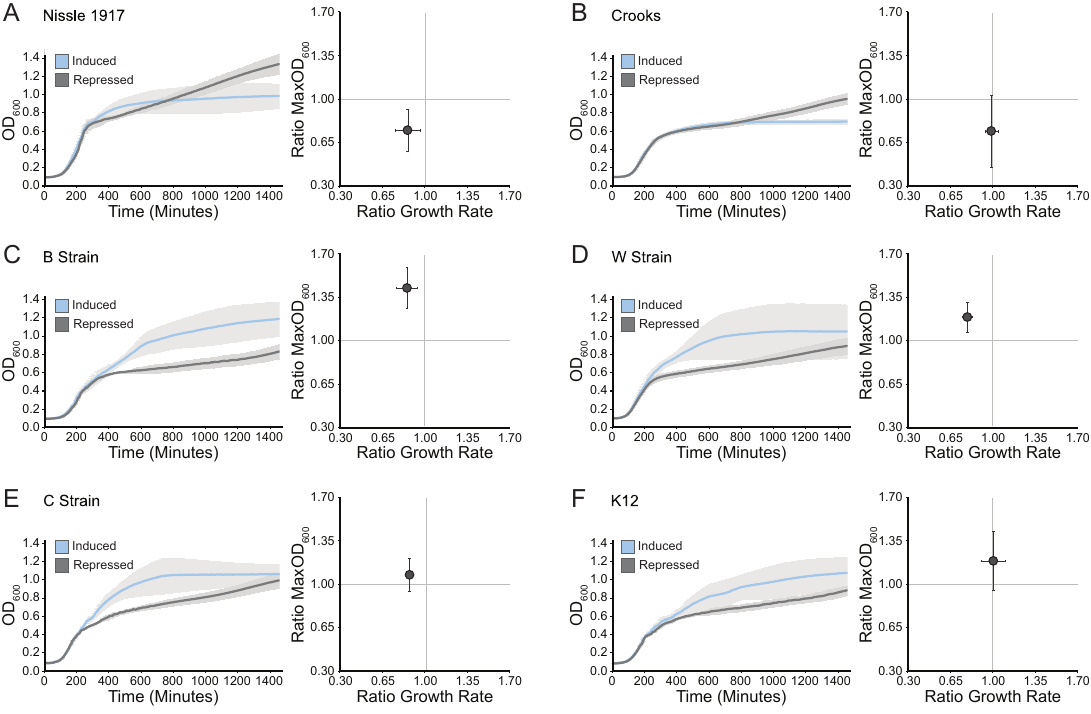

**Figure S10. Fitness analysis of six common laboratory strains expressing the mutant initiator tRNA-UUC.** Growth rate and maximum cell density were determined for strains grown in LB (spec) and induced with 1 mM IPTG, or repressed with 2% v/v glucose (n=3). **(Left)** Growth of cells carrying i-tRNA-UUC in an induced (blue) versus repressed (grey) condition. Line is the average value (n=3) and shading indicates bounds of one standard deviation. **(Right)** Growth rate and maximum cell density of cells expressing i-tRNA-UUC. Point represents average value n and error bars represent one standard deviation (n=3).

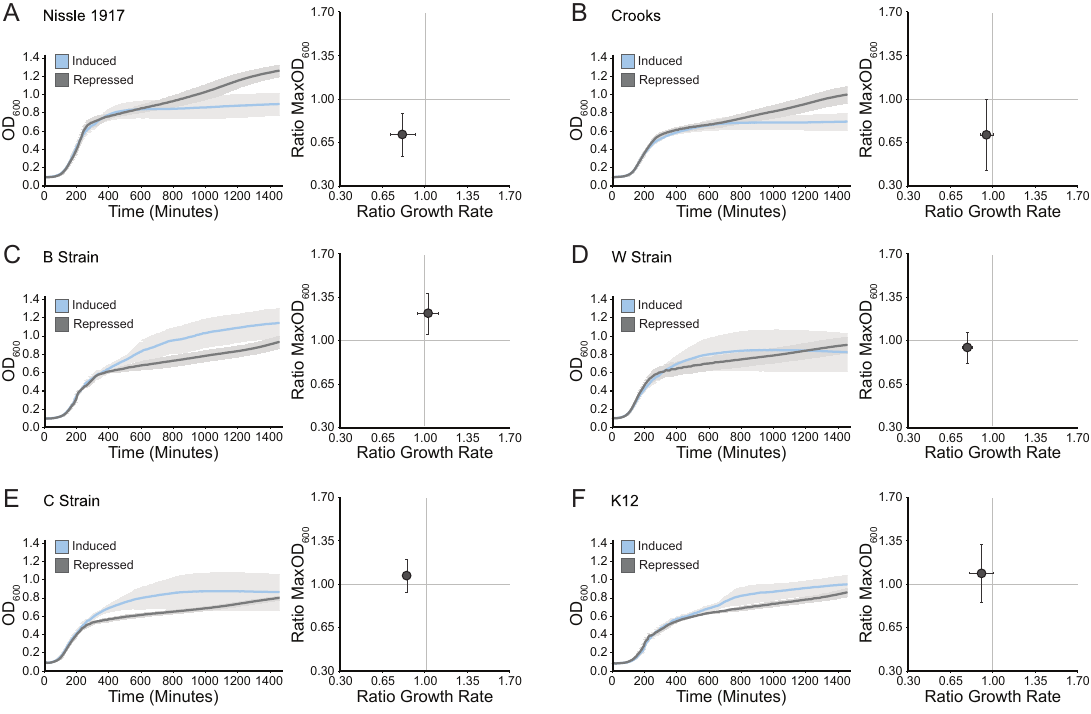
**Figure S11. Fitness analysis of six common laboratory strains expressing the mutant initiator tRNA-CUC.** Growth rate and maximum cell density were determined for strains grown in LB (spec) and induced with 1 mM IPTG, or repressed with 2% v/v glucose (n=3). **(Left)** Growth of cells carrying i-tRNA-CUC in an induced (blue) versus repressed (grey) condition. Line is the average value (n=3) and shading indicates bounds of one standard deviation. **(Right)** Growth rate and maximum cell density of cells expressing i-tRNA-CUC. Point represents average value n and error bars represent one standard deviation (n=3).

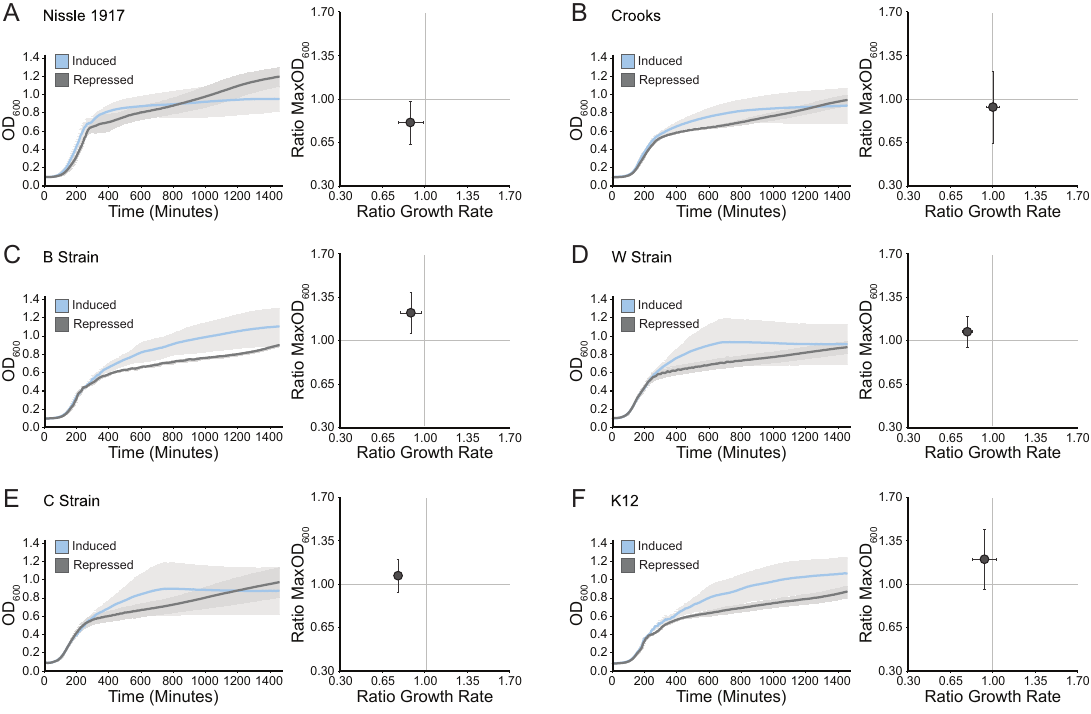

**Figure S12. Fitness analysis of six common laboratory strains expressing the mutant initiator tRNA-GGC.** Growth rate and maximum cell density were determined for strains grown in LB (spec) and induced with 1 mM IPTG, or repressed with 2% v/v glucose (n=3). **(Left)** Growth of cells carrying i-tRNA-GGC in an induced (blue) versus repressed (grey) condition. Line is the average value (n=3) and shading indicates bounds of one standard deviation. **(Right)** Growth rate and maximum cell density of cells expressing i-tRNA-GGC. Point represents average value n and error bars represent one standard deviation (n=3).

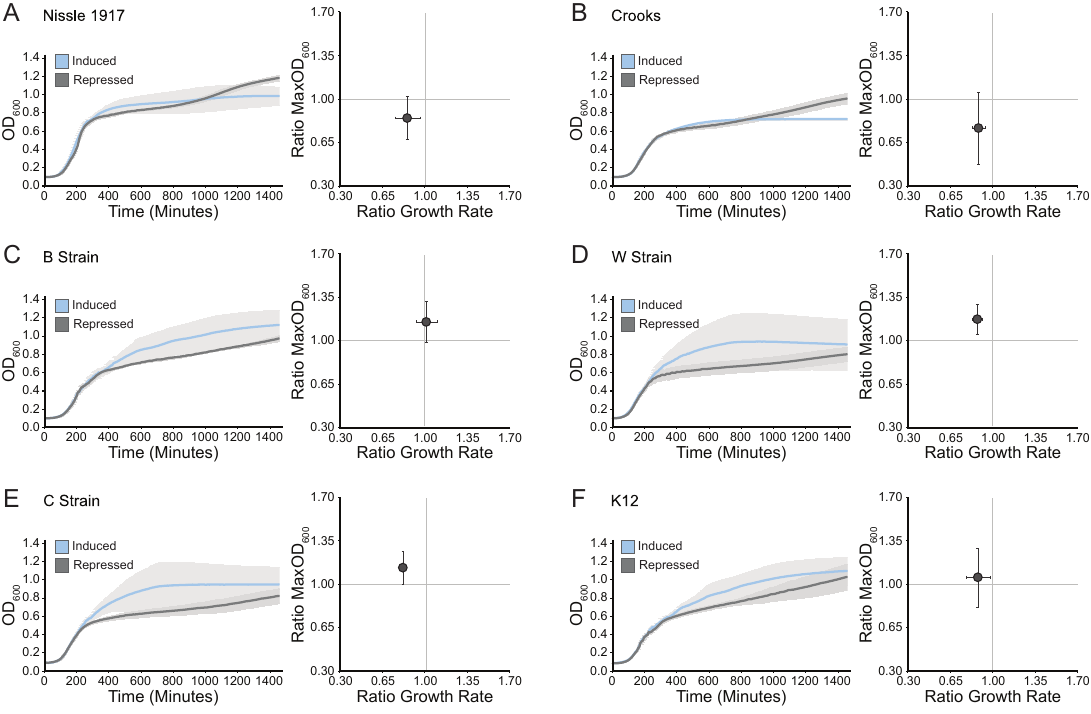
**Figure S13. Fitness analysis of six common laboratory strains expressing the mutant initiator tRNA-UAC.** Growth rate and maximum cell density were determined for strains grown in LB (spec) and induced with 1 mM IPTG, or repressed with 2% v/v glucose (n=3). **(Left)** Growth of cells carrying i-tRNA-UAC in an induced (blue) versus repressed (grey) condition. Line is the average value (n=3) and shading indicates bounds of one standard deviation. **(Right)** Growth rate and maximum cell density of cells expressing i-tRNA-UAC. Point represents average value n and error bars represent one standard deviation (n=3).

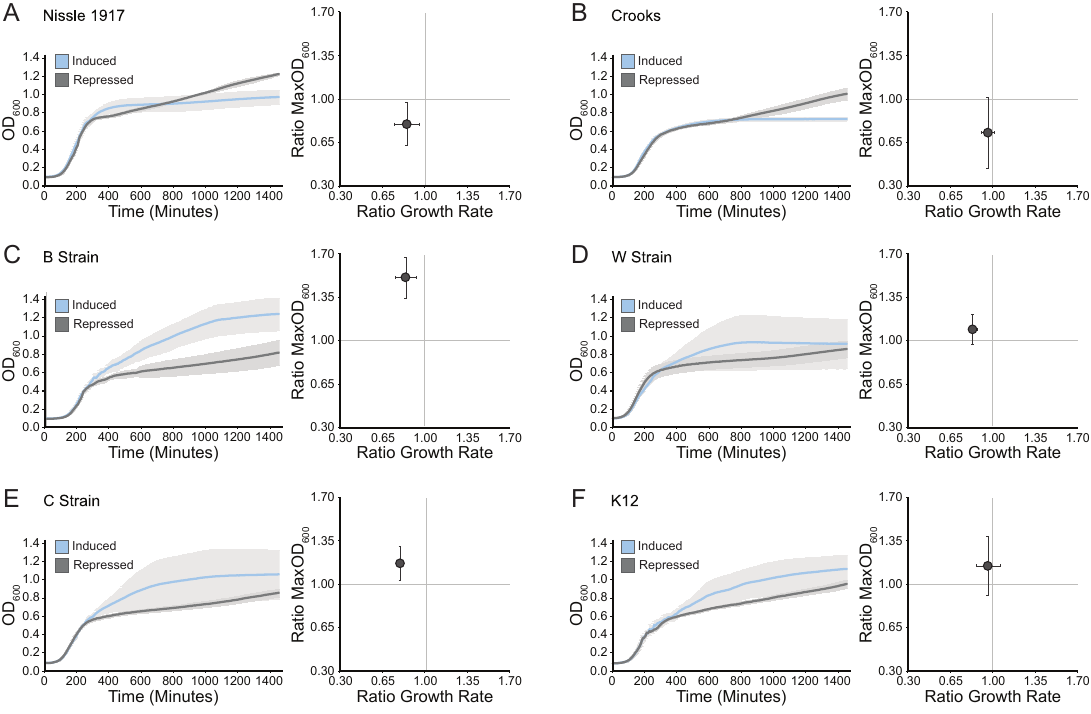

**Figure S14. Fitness analysis of six common laboratory strains expressing the mutant initiator tRNA-AAC.** Growth rate and maximum cell density were determined for strains grown in LB (spec) and induced with 1 mM IPTG, or repressed with 2% v/v glucose (n=3). **(Left)** Growth of cells carrying i-tRNA-AAC in an induced (blue) versus repressed (grey) condition. Line is the average value (n=3) and shading indicates bounds of one standard deviation. **(Right)** Growth rate and maximum cell density of cells expressing i-tRNA-AAC. Point represents average value n and error bars represent one standard deviation (n=3).

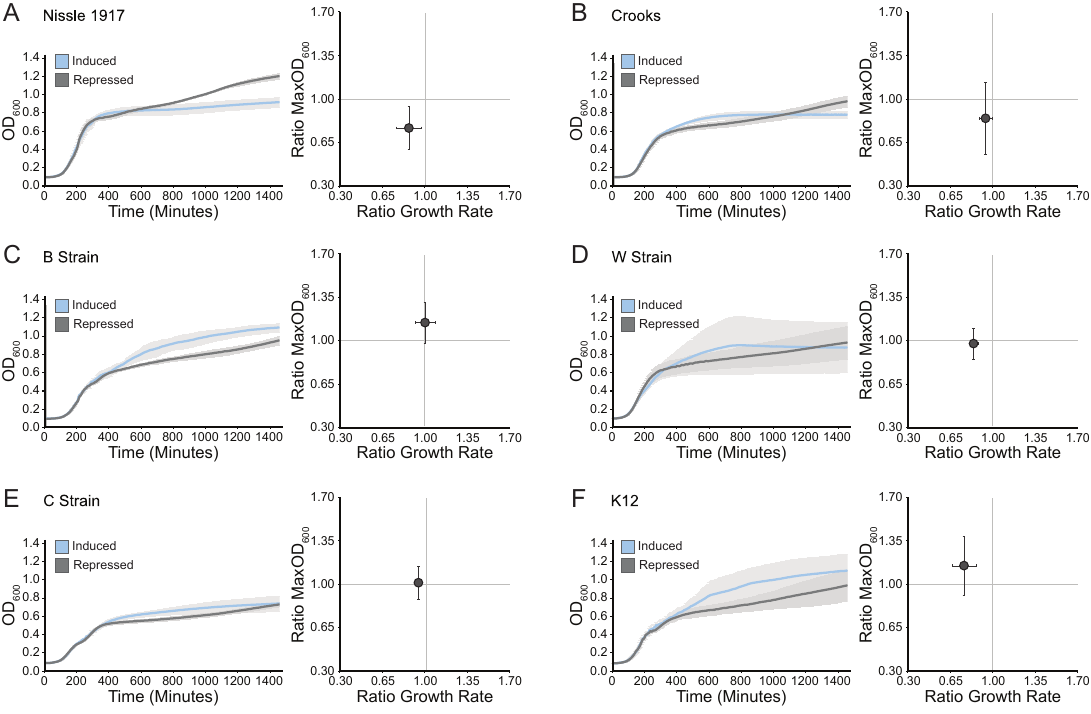
**Figure S15. Fitness analysis of six common laboratory strains expressing the mutant initiator tRNA-UUA.** Growth rate and maximum cell density were determined for strains grown in LB (spec) and induced with 1 mM IPTG, or repressed with 2% v/v glucose (n=3). **(Left)** Growth of cells carrying i-tRNA-UUA in an induced (blue) versus repressed (grey) condition. Line is the average value (n=3) and shading indicates bounds of one standard deviation. **(Right)** Growth rate and maximum cell density of cells expressing i-tRNA-UUA. Point represents average value n and error bars represent one standard deviation (n=3).

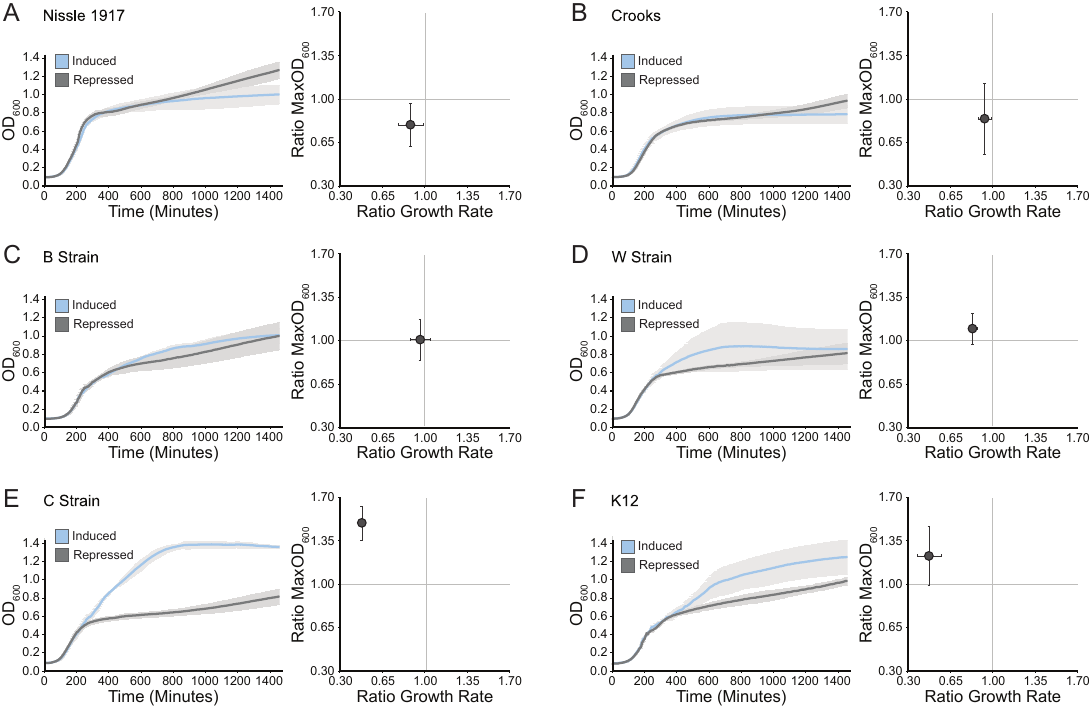

**Figure S16. Fitness analysis of six common laboratory strains expressing the mutant initiator tRNA-GUA.** Growth rate and maximum cell density were determined for strains grown in LB (spec) and induced with 1 mM IPTG, or repressed with 2% v/v glucose (n=3). **(Left)** Growth of cells carrying i-tRNA-GUA in an induced (blue) versus repressed (grey) condition. Line is the average value (n=3) and shading indicates bounds of one standard deviation. **(Right)** Growth rate and maximum cell density of cells expressing i-tRNA-GUA. Point represents average value n and error bars represent one standard deviation (n=3).

**Supporting Tables**

**Table S1. Two-way ANOVA of log_2_ fold-change in fluorescence**

| **term** | **df** | **sumsq** | **meansq** | **statistic** | **p.value** |
| --- | --- | --- | --- | --- | --- |
| **Strain** | 5 | 210.7133 | 42.14267 | 336.3271 | 1.59E-77 |
| **Codon** | 11 | 388.9601 | 35.36001 | 282.197 | 1.54E-91 |
| **Strain:Codon** | 55 | 309.7734 | 5.632244 | 44.94914 | 3.57E-68 |
| **Residuals** | 144 | 18.04357 | 0.125303 | NA | NA |

**Table S2. Two-way ANOVA of repressed-condition fluorescence**

| **term** | **df** | **sumsq** | **meansq** | **statistic** | **p.value** |
| --- | --- | --- | --- | --- | --- |
| **Strain** | 5 | 5.25321 | 1.050642 | 218.6022 | 2.21E-65 |
| **Codon** | 11 | 3.321315 | 0.301938 | 62.82279 | 2.44E-49 |
| **Strain:Codon** | 55 | 8.025467 | 0.145918 | 30.3604 | 5.13E-57 |
| **Residuals** | 144 | 0.69209 | 0.004806 | NA | NA |

**Table S3. Two-way ANOVA of induced-condition fluorescence**

| **term** | **df** | **sumsq** | **meansq** | **statistic** | **p.value** |
| --- | --- | --- | --- | --- | --- |
| **Strain** | 5 | 31.23344 | 6.246688 | 888.7403 | 1.18E-112 |
| **Codon** | 12 | 116.0252 | 9.668767 | 1375.613 | 1.62E-151 |
| **Strain:Codon** | 60 | 47.23394 | 0.787232 | 112.0026 | 3.58E-103 |
| **Residuals** | 156 | 1.096477 | 0.007029 | NA | NA |

**Table S4. Two-way ANOVA of growth rate**

| **term** | **df** | **sumsq** | **meansq** | **statistic** | **p.value** |
| --- | --- | --- | --- | --- | --- |
| **Strain** | 5 | 0.001959 | 0.000392 | 17.32256 | 2.11E-13 |
| **Codon** | 11 | 0.000664 | 6.04E-05 | 2.668958 | 0.003812 |
| **Strain:Codon** | 55 | 0.002371 | 4.31E-05 | 1.906432 | 0.001243 |
| **Residuals** | 144 | 0.003257 | 2.26E-05 | NA | NA |

**Table S5. Two-way ANOVA of maximum optical density**

| **term** | **df** | **sumsq** | **meansq** | **statistic** | **p.value** |
| --- | --- | --- | --- | --- | --- |
| **Strain** | 5 | 5.977854 | 1.195571 | 22.07792 | 2.23E-16 |
| **Codon** | 11 | 0.572563 | 0.052051 | 0.9612 | 0.484508 |
| **Strain:Codon** | 55 | 2.428627 | 0.044157 | 0.815419 | 0.805124 |
| **Residuals** | 144 | 7.797935 | 0.054152 | NA | NA |

**Table S6. Gene sequences.**

| **Gene** | **Sequence (5’-3’)** |
| --- | --- |
| *(NNN)sfgfp* | NNNCGTAAAGGCGAAGAGCTGTTCACTGGTGTCGTCCCTATTCTGGTGGAACTGGATGGTGATGTCAACGGTCATAAGTTTTCCGTGCGTGGCGAGGGTGAAGGTGACGCAACTAATGGTAAACTGACGCTGAAGTTCATCTGTACTACTGGTAAACTGCCGGTACCTTGGCCGACTCTGGTAACGACGCTGACTTATGGTGTTCAGTGCTTTGCTCGTTATCCGGACCATATGAAGCAGCATGACTTCTTCAAGTCCGCCATGCCGGAAGGCTATGTGCAGGAACGCACGATTTCCTTTAAGGATGACGGCACGTACAAAACGCGTGCGGAAGTGAAATTTGAAGGCGATACCCTGGTAAACCGCATTGAGCTGAAAGGCATTGACTTTAAAGAAGACGGCAATATCCTGGGCCATAAGCTGGAATACAATTTTAACAGCCACAATGTTTACATCACCGCCGATAAACAAAAAAATGGCATTAAAGCGAATTTTAAAATTCGCCACAACGTGGAGGATGGCAGCGTGCAGCTGGCTGATCACTACCAGCAAAACACTCCAATCGGTGATGGTCCTGTTCTGCTGCCAGACAATCACTATCTGAGCACGCAAAGCGTTCTGTCTAAAGATCCGAACGAGAAACGCGATCATATGGTTCTGCTGGAGTTCGTAACCGCAGCGGGCATCACGCATGGTATGGATGAACTGTACAAATGATGA |
| *metY(NNN)* | CGCGGGGTGGAGCAGCCTGGTAGCTCGTCGGGCTNNNAACCCGAAGATCGTCGGTTCAAATCCGGCCCCCGCAACCA |

**Table S7. Oligos used.**

| **Oligo** | **Sequence** |
| --- | --- |
| *metY* Sequencing and colony PCR Primer Forward | TCTCCCTTATGCGACTCCTG |
| *metY* Sequencing and colony PCR primer Reverse | AGATCCGGCCACGATGAC |
| *sfgfp* Mutagenesis Primer Forward | GAAATTAACCNNNCGTAAAGGCGAAG |
| *sfgfp* Mutagenesis Primer Reverse | TCCTCTTTAATGAATTCTGTGTGAAATTG |
| *sfgfp* colony PCR Primer Forward | CTCGCGTATCGGTGATTCAT |
| *sfgfp* colony PCR Primer Reverse | CGGATAACGAGCAAAGCACT |
| *sfgfp* Sequencing Primer Forward | CCTTTCGTCTTCACCTCGAG |
| *sfgfp* Sequencing PCR Primer Reverse | GTTACCAGAGTCGGCCAAG |
